# Exploiting Activation and Inactivation Mechanisms in Type I-C CRISPR-Cas3 for Genome Editing Applications

**DOI:** 10.1101/2023.08.05.552134

**Authors:** Chunyi Hu, Mason T. Myers, Xufei Zhou, Zhonggang hou, Macy L. Lozen, Yan Zhang, Ailong Ke

**Affiliations:** Department of Molecular Biology and Genetics, Cornell University, 253 Biotechnology Building, Ithaca, NY 14853, USA; Department of Biological Chemistry, University of Michigan, Ann Arbor, MI 48109, USA

**Keywords:** CRISPR, genome editing, Cascade, Cas3, anti-CRISPR, Acr, gene activation

## Abstract

Type I CRISPR-Cas systems utilize the RNA-guided Cascade complex to identify matching DNA targets, and the nuclease-helicase Cas3 to degrade them. Among seven subtypes, Type I-C is compact in size and highly active in creating large-sized genome deletions in human cells. Here we use four cryo-electron microscopy snapshots to define its RNA-guided DNA binding and cleavage mechanisms in high resolution. The non-target DNA strand (NTS) is accommodated by I-C Cascade in a continuous binding groove along the juxtaposed Cas11 subunits. Binding of Cas3 further traps a flexible bulge in NTS, enabling efficient NTS nicking. We identified two anti-CRISPR proteins AcrIC8 and AcrIC9, that strongly inhibit *N. lactamica* I-C function. Structural analysis showed that AcrIC8 inhibits PAM recognition through direct competition, whereas AcrIC9 achieves so through allosteric inhibition. Both Acrs potently inhibit I-C-mediated genome editing and transcriptional modulation in human cells, providing the first off-switches for controllable Type I CRISPR genome engineering.

## Introduction

CRISPR-Cas systems protect prokaryotes with RNA-guided adapted immunity (Barrangou et al., 2007; Brouns et al., 2008; Marraffini and Sontheimer, 2008). Short DNA fragments from the invader are integrated into the CRISPR locus as spacers to establish immunological memory (Barrangou et al., 2007; Yosef et al., 2012)). The transcript of the CRISPR array is processed into mature <u>CR</u>ISPR <u>RNAs</u> (crRNAs) to guide the Cas effector proteins to destroy the complementary nucleic acid target (Brouns et al., 2008; Garneau et al., 2010; Gasiunas et al., 2012; Hale et al., 2009; Jinek et al., 2012). CRISPR-Cas systems can be categorized into two major classes. Class 1 uses multi-subunit effector complexes for crRNA-guided target surveillance and destruction, whereas Class 2 uses a single-protein enzyme (Makarova and Koonin, 2015; Makarova et al., 2015). Many Class 2 effectors, such as Cas9 and Cas12, have been widely utilized for eukaryotic genome editing. Conversely, bacteriophages have evolved small inhibitor proteins known as the <u>a</u>nti-<u>CR</u>ISPRs (Acrs) to inactivate CRISPR effectors through diverse mechanisms (Bondy-Denomy et al., 2013; Marino et al., 2020). Acr proteins that thwart CRISPR-Cas9 have been exploited to control the temporal, spatial, or tissue distribution of gene editing activity, or to reduce undesired off-target events (Bubeck et al., 2018; Lee et al., 2019; Marino et al., 2020; Shin et al., 2017).

Class 1 CRISPR systems, including Types I, III, and IV, are far more abundant than Class 2 systems (Makarova et al., 2020). Type I alone accounts for nearly 50% of all CRISPR-Cas systems found in nature (Makarova et al., 2015). They are further divided into at least seven subtypes I-A through I-G (Makarova et al., 2020). They rely on a multi-subunit ribo<u>n</u>ucleo<u>p</u>rotein (RNP) complex called <u>C</u>RISPR-<u>as</u>sociated <u>c</u>omplex for <u>a</u>nti-viral <u>de</u>fense (Cascade) to identify a matching target site flanked by a 5’ <u>p</u>rotospacer-<u>a</u>djacent <u>m</u>otif (PAM), and a helicase-nuclease Cas3 subsequently recruited to full R-loop for processive DNA degradation (Brouns et al., 2008; Dillard et al., 2018; Hayes et al., 2016; Hochstrasser et al., 2014; Huo et al., 2014; Jackson et al., 2014; Mulepati and Bailey, 2013; Mulepati et al., 2014; Rutkauskas et al., 2015; Sashital et al., 2012; Sinkunas et al., 2013; Westra et al., 2012; Wiedenheft et al., 2011; Xiao et al., 2018; Xiao et al., 2017; Zhao et al., 2014). This stepwise activation model was found to hold true for most subtypes of Type I systems.

However, it was recently found that in I-A and I-G systems, Cas3 associates with Cascade constitutively to facilitate DNA target searching, and I-A Cas3 was found to be allosterically activated by Cascade upon R-loop formation on target DNA (Hu et al., 2022; Shangguan et al., 2022) On the application side, Type I CRISPRs have been used for genome manipulations in prokaryotes since 2014 (Zheng et al., 2020), and in eukaryotes since 2019 (Cameron et al., 2019; Dolan et al., 2019, Morisaka, 2019 #18; Pickar-Oliver et al., 2019). The intrinsic activity of Cascade-Cas3 was to introduce targeted large deletions, from a few hundred base pairs to as large as 200 Kb (Cameron et al., 2019; Dolan et al., 2019; Hu et al., 2022; Morisaka et al., 2019; Osakabe et al., 2020; Tan et al., 2022). Small <u>in</u>sertion and <u>del</u>etions (indels) have been created using a pair of FokI nuclease-fused Cascade (Cameron et al., 2019). Locus-specific transcriptome or chromatin modulation was achieved by using Cascade to block the movement of RNA polymerase, or fusing Cascade with transcriptional or epigenetic modulators (Chen et al., 2020; Pickar-Oliver et al., 2019; Young et al., 2019)).

I-F variants that lack Cas3 but encode <u>C</u>RISPR-<u>as</u>sociated transposase (CAST) systems have recently enabled targeted large DNA insertion into the human genome (Lampe et al., 2023). Deeper mechanistic and structural understandings of Type I CRISPR will likely inspire novel and safer gene-editing strategies.

Among seven subtypes, I-C is arguably the most streamlined Type I system. I-C Cascade consists of crRNA, Cas5, Cas7, Cas8, and Cas11; the latter is translated from a small <u>o</u>pen reading frame (ORF) hidden inside *cas8c* gene (McBride et al., 2020). Earlier structural work on *D. Vulgaris* (Dvu) and *B. halodurans* I-C system provided important insights for I-C Cascade assembly (Hochstrasser et al., 2016; Nam et al., 2012; O’Brien et al., 2020).

Recent work further defined the R-loop formation process in high-resolution (O’Brien et al., 2023). However, the RNA-guided DNA cleavage mechanism by I-C Cas3 remained elusive. We previously established a new I-C system from *N. lactamica* (Nla) that confers bacteria interference against plasmid (Tan et al., 2022). By including the “hidden” Cas11 component, we repurposed Nla I-C into a genome editing platform with favorable attributes such as high efficiency, smaller delivery payload, and ease of RNP production (Tan et al., 2022).

In this study, we elucidate the Cas3 nuclease activation mechanisms by comparing the cryo-EM reconstructions of the *Nla* I-C interference complexes in four different functional states. Further comparison with the well-characterized type I-E systems revealed the structural basis for their idiosyncratic mechanistic behaviors. For example, consistent with the recent study (O’Brien et al., 2023), the Cas11 subunits hold the non-target DNA (NTS) strand near the target DNA/guide RNA heteroduplex, rather than hiding the NTS behind themselves. We further define the structural basis for the R-loop-dependent Cas3 recruitment mechanism and the DNA substrate handover mechanism for the I-C system. The latter event was considered a critical mechanistic step in initiating DNA degradation, however, the experimental proof was indirect (Xiao et al., 2017). By capturing the pre-nicking state of Cascade-Cas3, we now provide direct structural evidence showing defining the entire NTS bulge being captured by Cas3. We further identified two highly efficient anti-CRISPR proteins against the Nla I-C system and revealed the structural basis for their mechanism of inhibition. AcrIC8 prevents Cascade from engaging DNA target through an allosteric mechanism, by preventing the Cascade from adopting the PAM-searching conformation; whereas AcrIC9 does so through competitive inhibition, by mimicking a PAM-containing dsDNA and directly occupying the PAM-recognition site in Cascade. Structural comparison with other I-C Cascade/Acr complexes (O’Brien et al., 2023) pinpoints to the source of the specificity and potency of these anti-CRISPR proteins. Importantly, in human genome editing experiments both Acrs potently inhibited I-C Cascade-Cas3 induced DNA deletion, as well as gene activation mediated by I-C Cascade fused with P65 transcription activator domain. They represent the first set of Acrs harnessed as off-switches for multi-subunit CRISPR gene editors, paving the way towards safer and better controlled Type I CRISPR applications.

## Results

### Reconstitution, biochemical and structural analysis of Nla I-C Cascade and Cas3

The Nla I-C Cascade RNP and Cas3 were recombinantly expressed and purified from *E. coli* as previously described (Tan et al., 2022) **(Figure S1A-G; Table S1)**. In the <u>e</u>lectrophoretic <u>m</u>obility <u>s</u>hift <u>a</u>ssays (EMSAs), NlaCascade bound the target DNA with an apparent Kd of ∼30 nM, whereas only non-specific interactions were detected at 100 nM for non-cognate DNA substrate **(Figure S1H)**. In the *in vitro* cleavage assay, addition of both Cascade and Cas3 to dual-labeled DNA target led to the selective nicking of NTS DNA at the PAM-proximal region of the R-loop. Further addition of ATP triggered processive DNA degradation. Both the target-<u>s</u>trand (TS) and NTS DNA were nicked at multiple locations at the PAM-proximal side **(Figure S2)**. These results are consistent with the uni-directional DNA deletion pattern in *ex vivo* human cell editing experiments by Nla I-C Cascade-Cas3 (Tan et al., 2022).

### Summary of the four functional states in NlaCascade-Cas3 captured by cryo-EM

To define the RNA-guided DNA cleavage mechanisms in high resolution, we determined a series of cryo-EM structures of I-C Nla Cascade and Cas3, each depicting a distinct functional state **(Figures 1, S3; Table S2)**. Nla Cascade in its resting state, without bound target DNA, was reconstructed from single particle cryo-EM at 3.7 Å resolution. I-C Cascade was found to share a stronger structural similarity to that of I-A Cascade, despite having a biochemical activity more similar to I-E and I-F Cascades. For example, both I-C and I-A Cascades have a longer helical backbone, and assembled from seven instead of six Cas7 subunits along the crRNA **(Figure 1D)**. These observations are consistent with the parallel work defining Dvu Type I-C Cascade structure and the R-loop formation mechanism (McBride et al., 2020; O’Brien et al., 2023; O’Brien et al., 2020).

**Figure 1.**
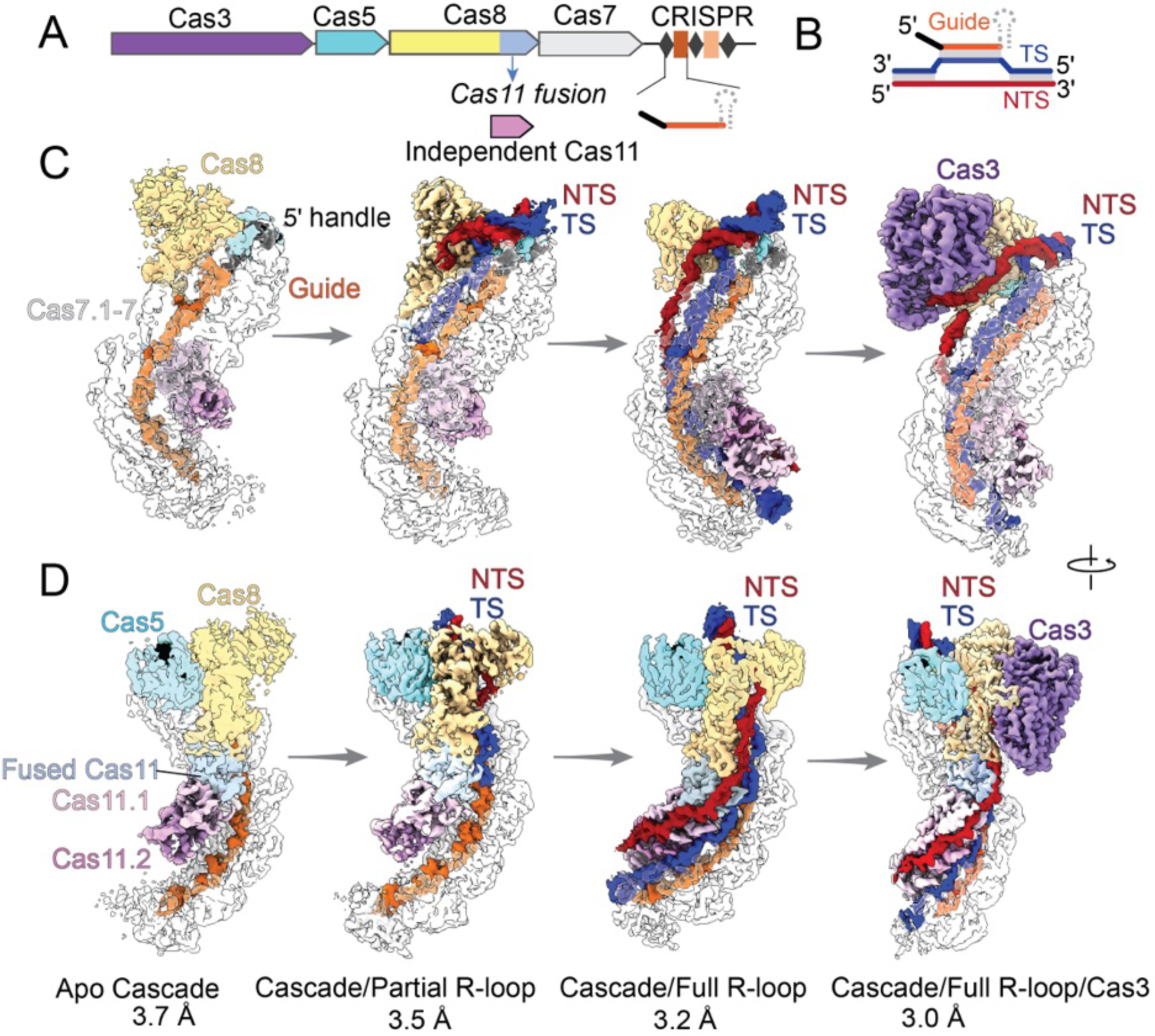
Four cryo-EM snapshots of *Nla* Type I-C CRISPR-Cas effector in different functional states. **(A)** Arrangement of the *Neisseria lactamica* Type I-C CRISPR-*cas* operon. **(B)** R-loop diagram between crRNA and target dsDNA. Coloring scheme is preserved throughout the figures: crRNA spacer (guide), orange; crRNA 5’ handle, black; target strand (TS) DNA, blue; non-target strand (NTS) DNA, red. **(C) and (D)** The overall cryo-EM maps at four different states, displayed in two different orientations. From left to right: the apo Cascade snapshot at 3.7 Å resolution; partial R-loop forming Cascade at 3.5 Å resolution; full R-loop forming Cascade at 3.2 Å resolution; and Cas3-bound Cascade/R-loop complex at 3.0 Å resolution.

Cryo-EM reconstruction resolved two functional states from the target DNA-bound Cascade sample **(Figures 1C-D)**. A 3.5 Å partial R-loop state was captured, in which the PAM-proximal 12 bp in the target DNA is unwound by I-C Cascade, and the target-strand in the unwound region is specified by crRNA through base-pairing interaction. The full R-loop structure was reconstructed at 3.2 Å resolution. 35 nucleotides of TS DNA are involved in base-pairing with the crRNA spacer, along the entire R-loop region. Structural comparison of these two states generated rich source of mechanistic insights. There are notable differences between how I-C and I-E Cascades accommodate the R-loop. Our previous work showed that I-E Cascade bends the target DNA by 45° at the PAM, and “hides” the NTS-DNA behind the oligomeric Cas11s, as much as 51 Å away from the TS-DNA/crRNA heteroduplex **(Figure 2A-B)** (Xiao et al., 2017). These structural features were interpreted as a mechanism to kinetically disfavor R-loop reversal (Xiao et al., 2017). Here, consistent with the recent observations from the Dvu I-C Cascade structures (O’Brien et al., 2023), Nla Cascade was found to only slightly bend dsDNA at the PAM region, by ∼10°, and uses a dedicated ssDNA-binding groove along Cas11 to accommodate the NTS. These sequence-nonspecific NTS binding units are located at the shoulder region of the oligomeric Cas11s, ∼20 Å above the TS-DNA/crRNA heteroduplex **(Fig. 2A, 2B)**.

**Figure 2.**
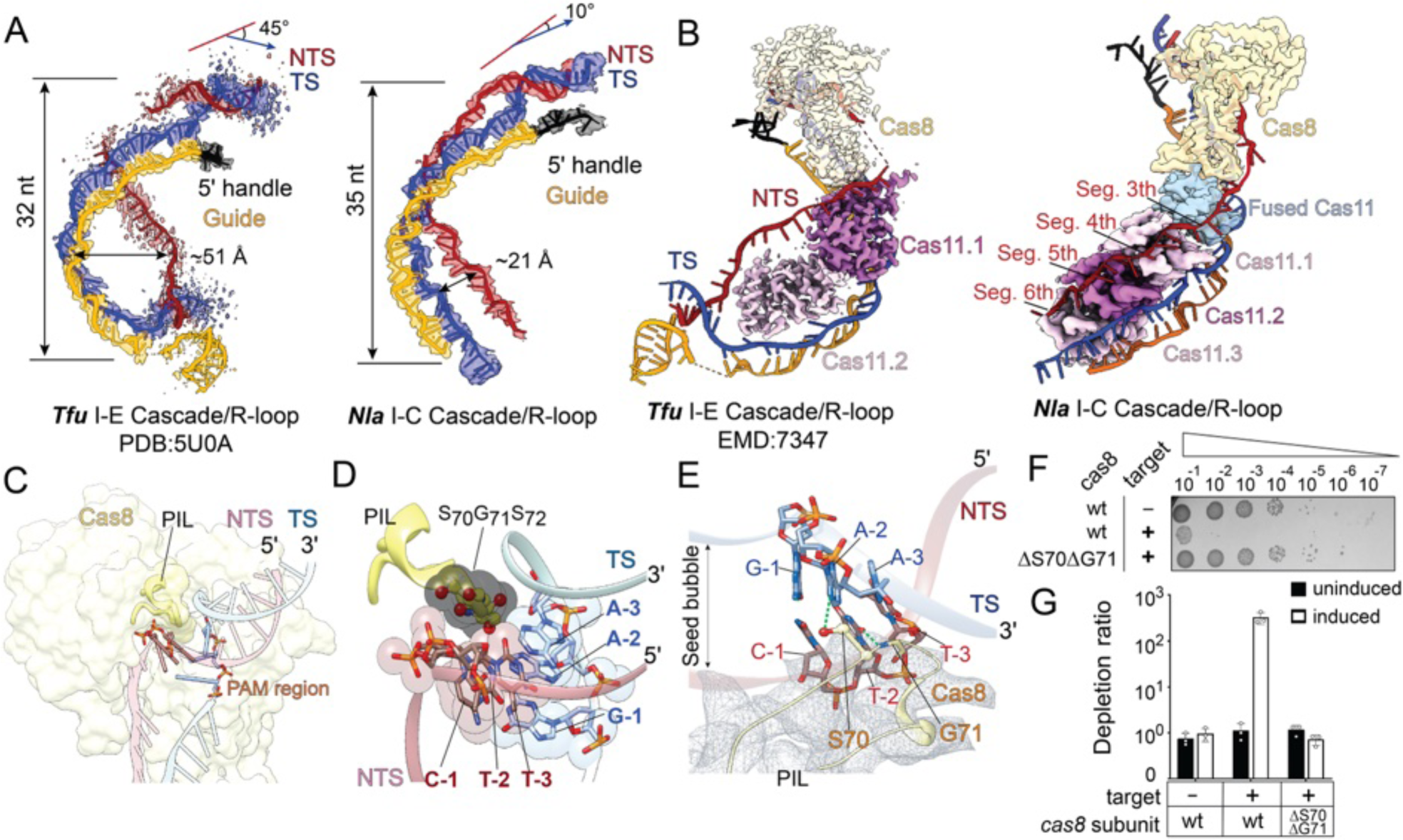
R-loop formation and PAM recognition mechanisms by *Nla* type I-C Cascade. **(A)** R-loop geometry inside *Tfu* Type I-E and *Nla* Type I-C Cascade, rendered in cryo-EM density and structural models. The NTS density inside I-E Cascade is poorly defined and contains a gap. The PAM-proximal dsDNA is significantly bent. **(B)** Differences in NTS accommodation in I-E and I-C Cascades. I-E Cascade ‘hides’ NTS to the backside of Cas11s. Contacts are limited to the NTS backbone. I-C Cascade has a dedicated NTS binding groove along Cas11 subunits. NTS is held not too far away from the TS-crRNA heteroduplex. **(C)** PAM recognition by Cas8 inside I-C Cascade. Note the PIL insertion into the minor groove of PAM, and the ss-NTS accommodation underneath. **(D)** PAM-T_-2_ and PAM-T_-3_ recognition from by the SGS-motif in PIL. Van de Waals spheres to show shape complementarity between S_70_G_71_S_72_ and PAM. (E) Another zoom-in view of the PAM contacts by the SGS-motif. **(F)** Importance of the SGS-motif evaluated using *E. coli*-based spot-titration based interference assay. ΔS_70_ΔG_71_ completely abolished the interference activity. Representative results are shown. **(G)** Same conclusion was reached from *E. coli*-based plasmid depletion assay. Depletion ratios were calculated as colony-forming units (CFUs) from the triple-antibiotic control plate divided by CFUs from the quadruple-antibiotic test plate for the same induced sample. Data are shown as mean ± SD, n = 3.

Nla I-C Cascade was further programmed to the Cas3-bound state for cryo-EM reconstruction. The resulting 3.0 Å structure depicts a pre-nicking state **(Figures 1C-D)**. Comparison of the structures before and after Cas3 recruitment allowed us to deduce the R-loop dependent Cas3 recruitment and activation mechanisms, which are explained in detail later.

### PAM recognition

Work in Type I-E and I-A systems (Hu et al., 2022; Xiao et al., 2017) showed that PAM recognition by Type I CRISPR systems typically involves Cas8-mediated DNA minor groove contacts. This typically coincides with the insertion of a wedge structure of Cas8 to initiate DNA duplex unwinding. These principles hold true for the Nla I-C Cascade, which specifies a 5’-T_-3_T_-2_Y(T/C)_-1_ PAM (Tan et al., 2022). In both the partial and full R-loop containing Cascade structures, a wedge structure in the <u>N</u>-terminal <u>d</u>omain (NTD) of Cas8 was found to unwind the DNA immediately underneath PAM. Above it, a hairpin loop in Cas8 inserts into the minor groove of the double-stranded PAM **(Figure 2C)**. This <u>P</u>AM-interaction loop (PIL) leans towards the NTS of the PAM duplex, due to several favorable sugar phosphate contacts from PIL and surrounding structures to NTS, including R69 to the phosphate of A_1_, N92 and K95 to the phosphate of C_-1_, and main chain G71 to T_-3_. A S_70_G_71_S_72_ (SGS) motif at the tip of PIL presses against T_-3_ and T_-2_ from the minor groove side. Despite the presence of heteroatoms, we think this SGS motif specifies PAM through Van der Waals contacts rather than hydrogen bonds **(Figures 2D-E)**. The nucleobases at PAM-2 and PAM-3 are specified as thymines through two levels of negative selections. First, the SGS motif only tolerates pyrimidines in the NTS PAM because the bulker purines cause steric clashes. Second, cytosines are rejected from these two positions because the N2 exocyclic amines in the compensatory guanines would clash with the SGS motif. In the Dvu I-C Cascade structure, The TT-PAM are recognized by a GNG motif (O’Brien et al., 2023). The molecular recognitions are compatible with our interpretation above. These structural observations recapitulate the recognition-through-elimination based PAM recognition mechanism first defined in the I-E Cascade structure (Hayes et al., 2016). Y_-1_ specification cannot be fully explained by the current structure. The potential hydrogen bonds between the Cas8 wedge and G_-1_ are not optimal in bond angle. It is possible that PAM-1 is specified during PAM searching, by residues in the wedge loop of Cas8. The subsequent DNA unwinding step may have shifted the local geometry and distorted the PAM-1 contact.

Consistent with our structural observations, disrupting the SGS motif in *cas8* (ΔS70ΔG71) completely abolished the plasmid interference function of Nla I-C *in vivo* **(Figures 2F-G)**, validating its functional importance. Moreover, this PAM recognition motif is highly conserved among many Cas8c homologs **(Figure S4A)**, suggesting that it is a good predictor of PAM specificity for I-C systems.

### Cas3 recruitment through handshaking with Cascade and NTS bulge capture

Nla I-C Cas3 only co-purified with Cascade in the presence of the cognate DNA substrate (data not shown), after which the DNA was selectively degraded **(Figure S2)**. Here we explain this trans-recruitment mechanism by comparing the Cas3-bound structure with the Cascade-only structures in various states. Several conformational changes occur when Cascade transitions from partial to the full R-loop state. A global conformational change takes place, which involves all subunits in the inner belly of Cascade. However, these components move to various extents. Whereas the Cas8 NTD (AA 1-259) slides on average 7-8 Å, the Cas11 subunits and Cas8 C-terminal domain (CTD, AA 458-582) only twist in place. The mid-region of Cas8, including the wedge domain (AA 260-348) and the adaptor domain (AA 349-457), undergoes a mixed range of twisting and sliding motions **(Figure S4B)**. The non-uniform conformational changes alter the Cas8 surface significantly. Moreover, a previously unstructured loop (A276-P297) in the wedge domain becomes well-ordered. It extends from the Cas8 surface, like a distinct welcoming hand (the recruit loop, **Figure S4C)**.

The Cas3-bound ternary structure further explains how Cas3 selectively binds to the Cascade/full R-loop through a conformation-capture type of mechanism **(Figure 3).** Cas3 HD nuclease domain interacts with the wedge domain of Cas8, and Cas3 helicase domain interacts with the Cas8 NTD **(Figure 3A, S4B)**. Because as explained earlier these two Cas8 domains undergo uneven conformational stretching upon full R-loop formation, the same set of Cas3-Cas8 contact would not be possible in the resting or partial R-loop states. Importantly, Cas3 binding is further strengthened by two reciprocal hand-shaking interactions: one from the recruitment loop of Cascade and the other from the anchor loop of Cas3 **(Figures 3A-C)**. The Cas8 recruitment loop only reaches up in the full R-loop state. It contacts Cas3 HD domain through an extensive set of hydrophobic interactions **(Figure 3B)**; the interface residues on both sides are well conserved among I-C systems **(Figures 3B, S4D)**. Conversely, the anchor loop from Cas3 reaches down into a groove in Cas8 NTD, which is only exposed in the full R-loop state. Its backbone is contacted by polar residues surrounding the Cas8 groove, and its conserved hydrophobic tip (F437) is pressed into a conserved hydrophobic pocket in Cas8 **(Figures 3C, S4E)**.

**Figure 3.**
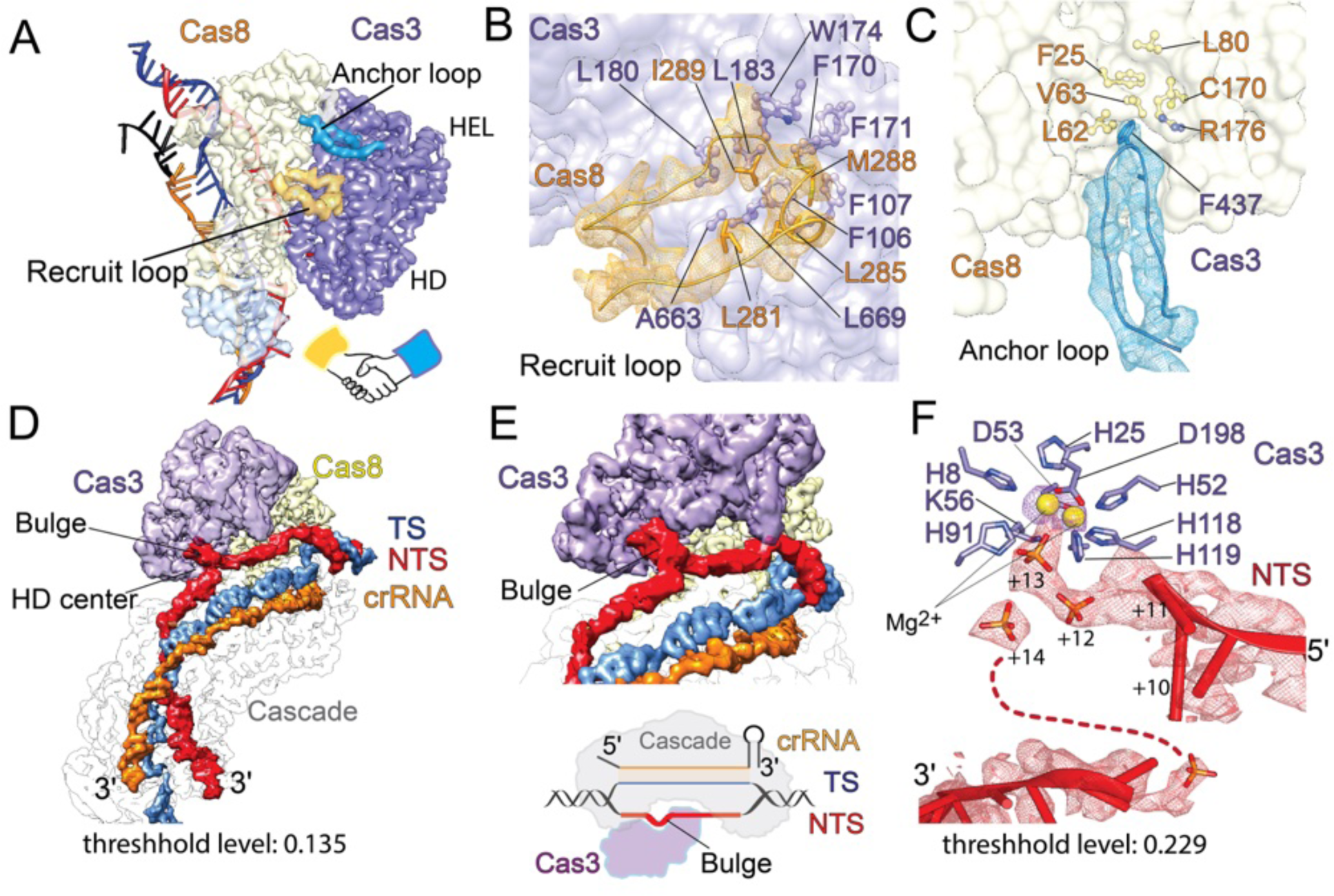
The activation mechanism of type I-C CRISPR-Cas3. **(A)** Cryo-EM densities showing that the R-loop forming Cascade recruits Cas3 through a hand-shaking mechanism. The blue anchor loop from Cas3 and the orange recruit loop from Cas8 reach out to mediate contacts. **(B)** Detailed interactions between the recruitment loop of Cas3 (density in yellow mesh) and its hydrophobic binding pocket in Cas8. **(C)** Detailed interactions between the anchor loop of Cas8 (density in cyan mesh) and its hydrophobic binding pocket in Cas3. **(D)** and **(E)** Overall and zoom-in views of the cryo-EM reconstruction showing the NTS bulging up from its regular path, reaching into the Cas3 HD nuclease center, then making a 180° turn to resume its regular path on the Cascade surface. **(F)** A more close-up view of the NTS bulge in and out of Cas3 HD center, with cryo-EM densities contoured at a higher level. Mobile regions coincide with choppy densities.

Upon binding, the HD nuclease center of Cas3 is positioned ∼20 Å above its substrate, the NTS ssDNA. Previously, we predicted that R-loop formation traps a flexible NTS ssDNA bulge, which is then captured and nicked by the nearby Cas3 HD nuclease. This mechanism was inferred from the structural work in I-E system revealing that the NTS density is disordered in this area, and some residual densities hint the handover mechanism (Xiao et al., 2018). To our surprise, we only observed continuous NTS densities in the full R-loop Cascade structure; there was no indication of DNA bulge formation at the expected Cas3 binding surface **(Figure 2A-B)**. This puts the NTS handover model in question. This discrepancy was resolved in the Cas3-bound I-C Cascade/R-loop structure, which not only captured the NTS bulge, but also revealed its entire path **(Figure 3D-E)**. The NTS DNA was found to rise sharply from its surface path at NTS_+11_, enters Cas3 HD active site at NTS_+13_, makes a sharp U-turn afterwards and joins the original path at NTS_+16_ **(Figure 3E)**. The EM density following NTS_+13_ is weaker and more degenerate, which could be due to low levels of NTS nicking by Cas3. Our structures suggest that slack is present and dynamically distributed along the NTS upon Cascade-mediated R-loop formation. It becomes concentrated upon Cas3 binding, which enables Cascade to capture NTS for strand-nicking.

### *In vivo* functional assay defines two Acrs inhibitors against Nla I-C CRISPR-Cas

We sought to identify Nla I-C specific Acrs that can potentially serve as off-switches in human cell genome editing experiments. Previous studies have predicted nearly a dozen potential Type I-C-specific Acrs based on gilt-by-association, self-targeting, and machine learning approaches; they were shown to block *<u>P</u>. <u>ae</u>ruginosa* (Pae) I-C CRISPR-Cas in phage defense (Gussow et al., 2020; Leon et al., 2021). However, because Acrs typically establish highly specific physical interactions with their corresponding Cas effectors, their efficacy on other given I-C systems could vary significantly. We therefore performed plasmid interference assays in *E coli* to screen for efficient Acrs against our Nla I-C system. Among twelve candidates tested, Acrs IC8 and IC9 were found to be potent inhibitors **(Figures 4A-B, S5A-B)**. Next, we turned to a DNA-binding based GFP repression assay **(Figure 4C)** to discern if they inhibit Nla I-C at the step of DNA binding by Cascade or later steps. *E. coli* BW25113 derivative strains expressing Cascade, a LacZ promoter driven GFP reporter, and an arabinose inducible Acr **(Figure S5C)** were assayed for GFP fluorescence. Functional Nla Cascade is guided by its crRNA to bind a target site adjacent to the *lacI* promoter and thereby blocks *lacI* transcription. This in turn alleviates the *lacI*-mediated repression of GFP expression, leading to green, fluorescent cells **(Figure 4C)**. Induction of AcrIC8 or AcrIC9 expression led to a 5.4-or 14.3-fold reduction in GFP fluorescence, respectively, whereas the no Acr or *S. pyogenes* (Spy) Cas9-specific AcrIIA4 (Rauch et al., 2017) negative controls only caused modest ∼2-fold GFP repressions (**Figures 4D-E**). Our results showed that among the many AcrICs shown to be effective against *P. aeruginosa* I-C Cascade (Leon et al., 2021), only AcrIC8 are effective against Nla Cascade. Similarly, AcrIC4 was found to strongly inhibit Dvu I-C Cascade, however, it had no effect on Nla Cascade (O’Brien et al., 2023). It is therefore important to define the suitable anti-CRISPRs for a given CIRSPR system through systematic functional screens.

**Figure 4.**
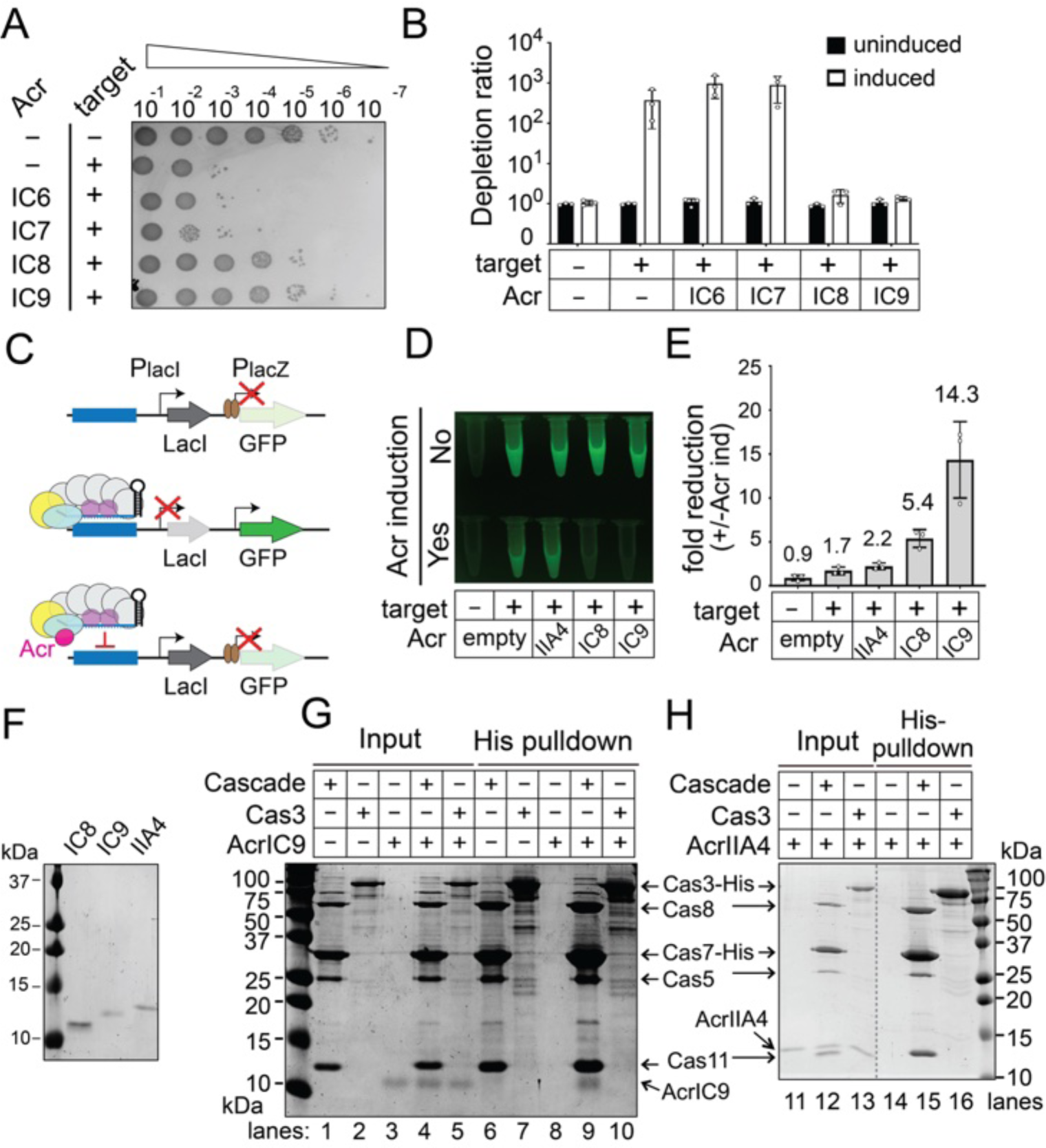
AcrIC8 and AcrIC9 inhibits Nla I-C CRISPR function by preventing DNA binding. **(A-B)** Acrs IC8 and IC9 disrupt Nla I-C CRISPR interference *in vivo*. A representative image of titration-spotting *E. coli* interference assay was shown in (A). Colony reduction caused by Nla I-C CRISPR-Cas was unaffected by Acrs IC6 or IC7 but abrogated by IC8 and IC9. Depletion ratios are quantified and plotted in (B) as in Figure 2E. **(C)** Schematics of the CRISPR-i based transcription repression assay. Top, GFP reporter expression is blocked by LacI repressors (brown circles) produced from the upstream LacI cassette. Middle, Nla Cascade binding to a target site present in lacI promoter leads to transcriptional blockage of LacI and in turn activates GFP expression. Bottom, an Acr that disrupts Cascade binding to target DNA would restore LacI production and GFP repression. **(D)** GFP fluorescence image of OD_600_ normalized *E. coli* cultures from a representative CRISPR repression experiment. Fold of GFP repression by Acr was quantified by microplate reader and plotted in **(E)**. For each strain, repression was calculated as the fluorescence of uninduced sample normalized against its OD_600_ value, divided by OD_600_-normalized fluorescence of the induced sample. Data shown are mean ± SD, n = 3. **(F)** 15% SDS-PAGE showing the purity of AcrIC8, IC9, and IIA4 used in this work. **(G)** His-tag pulldown experiment revealing direct interactions between untagged AcrIC9 and His_6_-tagged Nla Cascade. The input and pulldown fractions are shown on the left and right portion of the Coomassie-stained SDS-PAGE. **(H)** Negative control experiment done in the same setup as in (G) showing the lack of interaction between SpyCas9-specific AcrIIA4 and Nla Cascade.

Next, we conducted *in vitro* nickel affinity pull-down experiments to test if these two Acrs physically interact with Nla Cascade. Under stringent conditions, the untagged Acr (**Figures 4F, S5D-K)** only co-elutes with the His_6_-tagged bait if they form a complex. Results showed that AcrIC9 was pulled down by His_6_-Cascade but not His_6_-Cas3, suggesting that it specifically targets Cascade **(Figure 4G)**. By contrast, the SpyCas9-specific AcrIIA4 did not co-elute which Cascade or Cas3 **(Figure 4H)**. Of note, untagged AcrIC8 alone can be pulled down by nickel resin non-specifically, precluding any conclusion about its association with Cascade or Cas3 *in vitro* (**Figure S5L**).

### AcrIC8 prevents DNA binding by Cascade through allosteric inhibition

We reconstituted the AcrIC8- and AcrIC9-bound Cascade complexes and determined their cryo-EM structure at 3.4 and 3.6 Å resolution, respectively (**Figures 5, 6 and S6**). While both Acrs bind to the PAM-proximal region of Cascade, they prevent DNA binding through distinct mechanisms. AcrIC8 inactivates Cascade by trapping the PAM-recognizing Cas8 subunit in a non-productive conformation, incapable of performing PAM recognition. The flat-shaped AcrIC8 wedges deep into the TS-DNA binding grove and anchors itself to the first segment of crRNA spacer **(Figures 5A-B)**. Besides the overall charge and shape complementarities at the binding interface, the following contacts contribute to AcrIC8’s specific efficacy against Nla I-C Cascade. A short loop of AcrIC8 (G21-E29) probes a Cascade cavity between the wedge and adaptor domains of Cas8 **(Figures 5C-D)**, forming a network of favorable hydrogen bonds (E22, E23 of AcrIC8 to R265, K389 of Cas8; **Figure 5E**). A separate set of specific contacts is found at the bottom of the crRNA displaying groove (**Figure 5E**). Y11 of AcrIC8 packs against a hydrophobic patch on Cas7.2 (L73 and V115), forming a specific hydrogen bond with R149 in the thumb domain of Cas7.1 (**Figure 5E**). Nearby, R58 of AcrIC8 mediates bidentate hydrogen bonds with D163 and D157 of Cas7.1, and S56 of AcrIC8 mediates hydrogen bonds with N154 and K156 of Cas7.1 (**Figure 5E**). Y11, R58, and E22 are absolutely conserved among AcrIC8 homologs; E23 and S56 are conserved in a subgroup **(Figure S7A)**.

**Figure 5.**
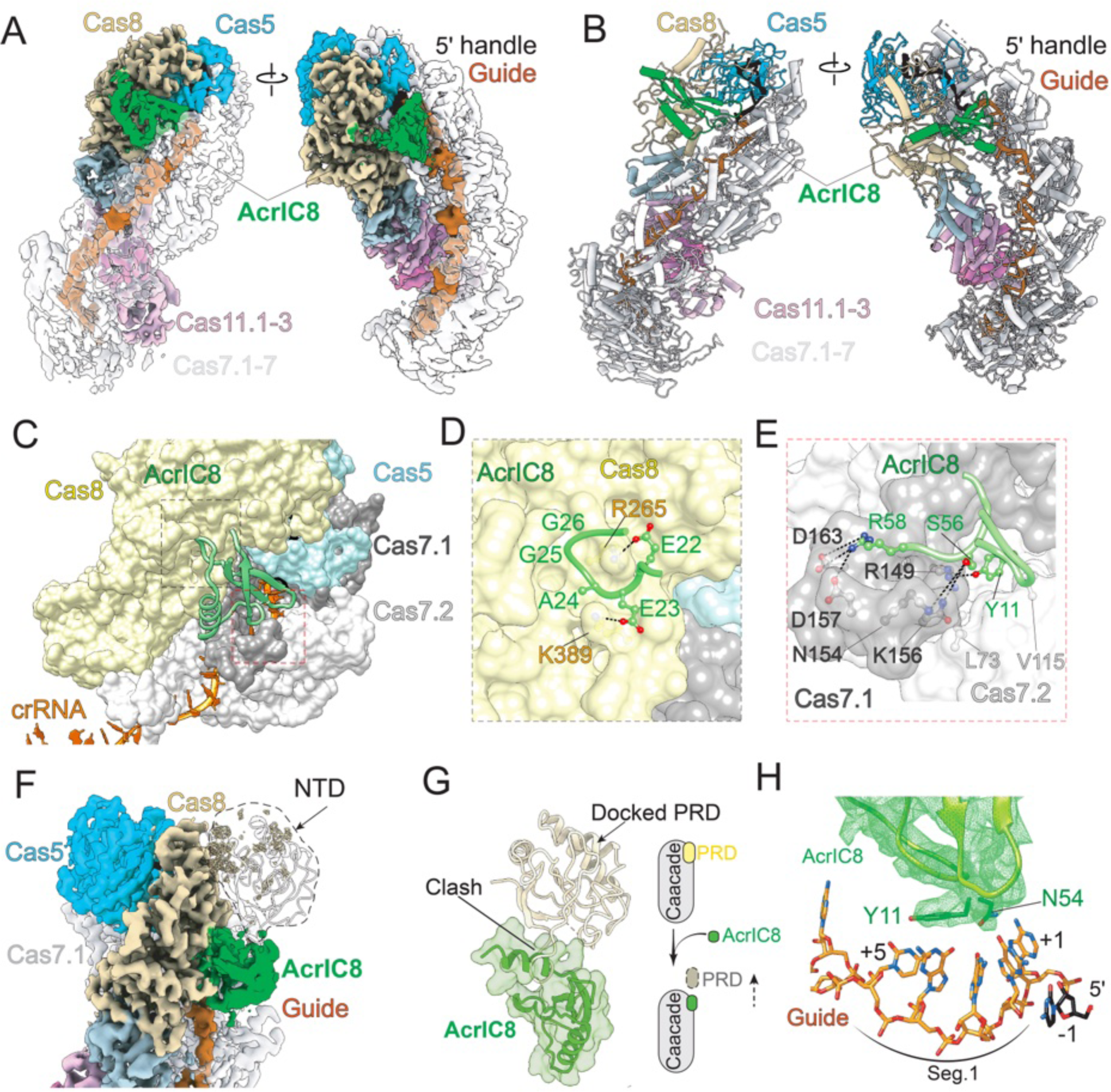
Structural basis for AcrIC8-mediated Nla I-C Cascade inactivation. **(A)** Overall cryo-EM density and **(B)** structural model of AcrIC8-bound Nla I-C Cascade. **(C)** The close-up view of the AcrIC8-Cascade interface. AcrIC8 contacts to Cas8 and Cas7.1 subunits are boxed and are displayed in details in panels **(D)** and **(E)**, respectively. **(F)** NTD of Cas8 disappeared from the cryo-EM density upon AcrIC8 binding. **(G)** Superposition of the apo and AcrIC8-bound Cascade structure reveals a steric clash between Cas8 NTD and AcrIC8. This leads to the mechanistic interpretation that AcrIC8 inhibits Cascade allosterically, by trapping Cas8 in the inactive conformation. **(H)** AcrIC8 further blocks the Watson-Crick edge of crRNA, preventing base-pairing formation with the target DNA.

While AcrIC8 does not directly compete for dsDNA binding, it prevents Nla Cascade from doing so through allosteric controls. First its binding sterically precludes the NTD of Cas8 **(Figure 5F-G)** from adopting the PAM-recognition conformation seen in Cascade structures **(Figure 1)**; Cas8 NTD becomes highly mobile and disappears from the EM density **(Figures 5A and 5F)**. Secondly, by occupying where the first segment of TS DNA binds on crRNA, AcrIC8 prevents the Cascade from specifying a DNA target (**Figure 5H**).

### AcrIC9 competes with dsDNA from Cascade by mimicking PAM

Our structure analysis revealed that AcrIC9 directly compete for the PAM recognition side in Nla Cascade **(Figures 6A-C)**. The 84-AA AcrIC9 mimics a double-stranded PAM in shape and charge distribution. It assumes the similar width as a dsDNA, and further displays two stripes of negative charges (E63/E64/D68 on one side and Q13/E54/D57/D58 on the other) to mimic the antiparallel sugarphosphate backbones (**Figures 6C-E**). The structural features in AcrIC9 tricks the Cascade to make the same set of PAM contacts **(Figures 6F-H and S7B)**. The two stripes of negative charges are accommodated by the same structural features in Cas8 that contact the sugarphosphate backbones of the double-stranded PAM **(Figure S7B-C)**. The SGS motif in the PIL of Cas8 probes into the groove in the middle, making multiple Van der Waals contacts to a hydrophobic patch on AcrIC9 consisting of F4, W18, V59, and W67 (**Figure 6G**). Like AcrIC8, AcrIC9 also uses a highly conserved tyrosine residue Y10 to probe the interface between Cas7.1 and Cas7.2 (**Figure 6H**). The aforementioned interface residues in AcrIC9 are well conserved **(Figure S7D)**, suggesting that all AcrIC9 homologs disables I-C Cascade through PAM mimicry.

**Figure 6.**
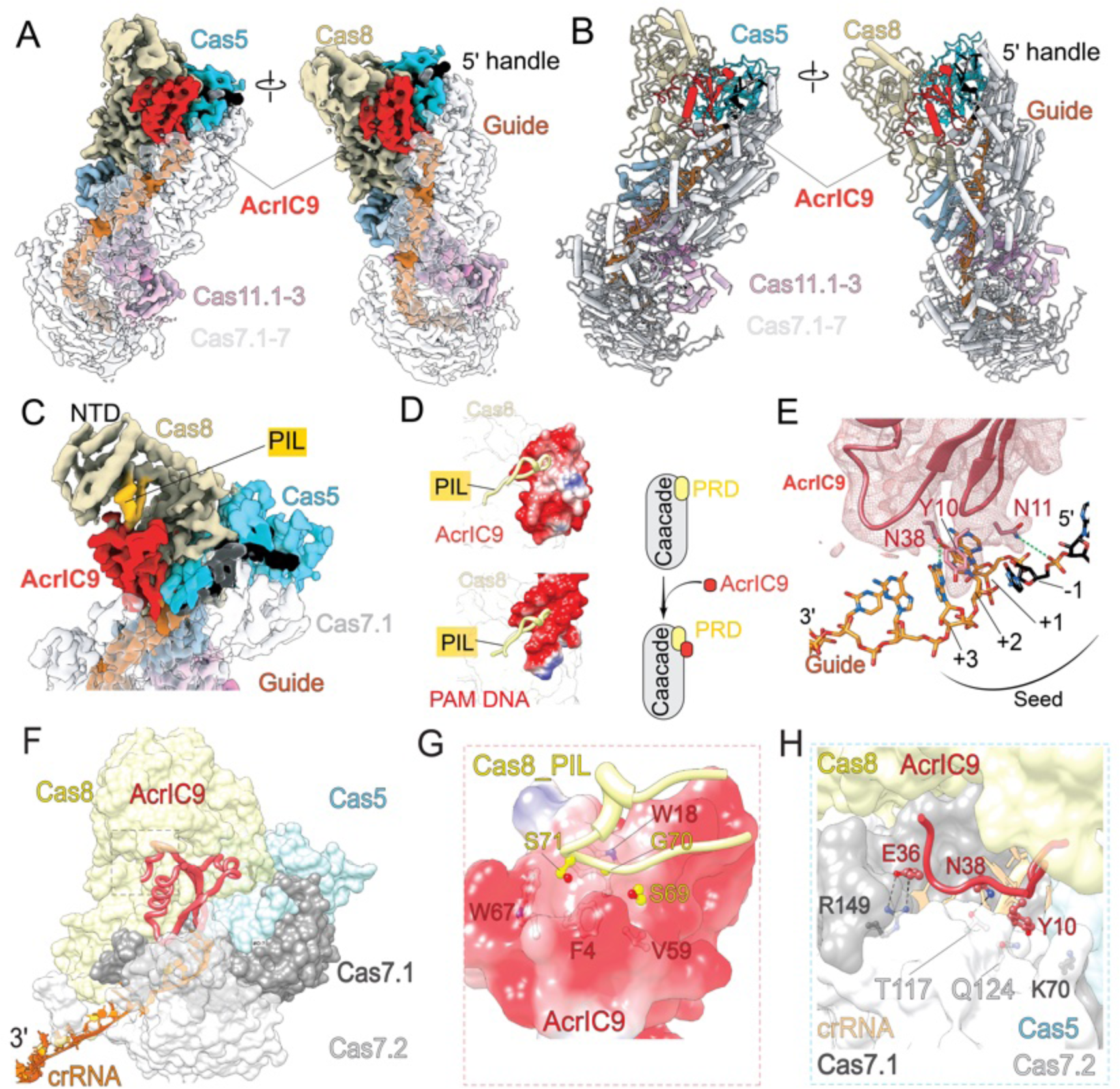
Structural basis for AcrIC9-mediated Nla I-C Cascade inactivation. **(A)** Overall cryo-EM density and **(B)** structural model of AcrIC9-bound Nla I-C Cascade. **(C)** Close-up view showing that AcrIC9 occupies the PAM binding site in Cas8, sequestering the PAM interaction loop. **(D)** AcrIC9 mimics dsDNA in shape and electrostatic distribution. PIL-PAM interaction is sown at the bottom for comparison. **(E)** AcrIC9 further blocks the Watson-Crick edge of crRNA, interfering with target DNA specification. **(F)** Orientation view showing the position of AcrIC9 inside Cascade. **(G)** PIL-AcrIC9 contacts are hydrophobic in principle, which are similar to the minor groove PAM contacts by PIL. **(H)** Specific contacts from AcrIC9 to Cas7.1 and Cas7.2. Collectively, these contacts (D-H) enabled AcrIC9 to occupy the PAM binding site with high affinity, preventing dsDNA binding through competitive inhibition.

### AcrIC8 and AcrIC9 as potent inhibitors in mammalian gene editing

Next, we evaluated the potency of AcrIC8 and AcrIC9 in serving as off-switches for Nla I-C gene editor in cultured human cells. We co-transfected HAP1-GFP reporter cells with a mixture of plasmids expressing Cas3, a GFP-targeting Cascade, and a Acr protein. Gene targeting/deletion efficiencies were measured four days later by flow cytometry (**Figure. 7B**). We observed ∼30% GFP disruption by NlaCascade-Cas3 when no Acr or AcrIE2, a negative control Acr, was included. AcrIC8 and AcrIC9 each greatly repressed gene editing in a dose-dependent fashion. Plasmid titration showed that at the lowest amount (20 ng) used, AcrIC8 and AcrIC9 reduced editing efficiency to 7.7% and 1.4%, respectively (**Figure. 7C**). When used at higher amounts (100-150 ng), both Acrs exhibited near-complete inhibition of genome editing by Nla I-C system to the background level (**Figure. 7C**).

**Figure 7.**
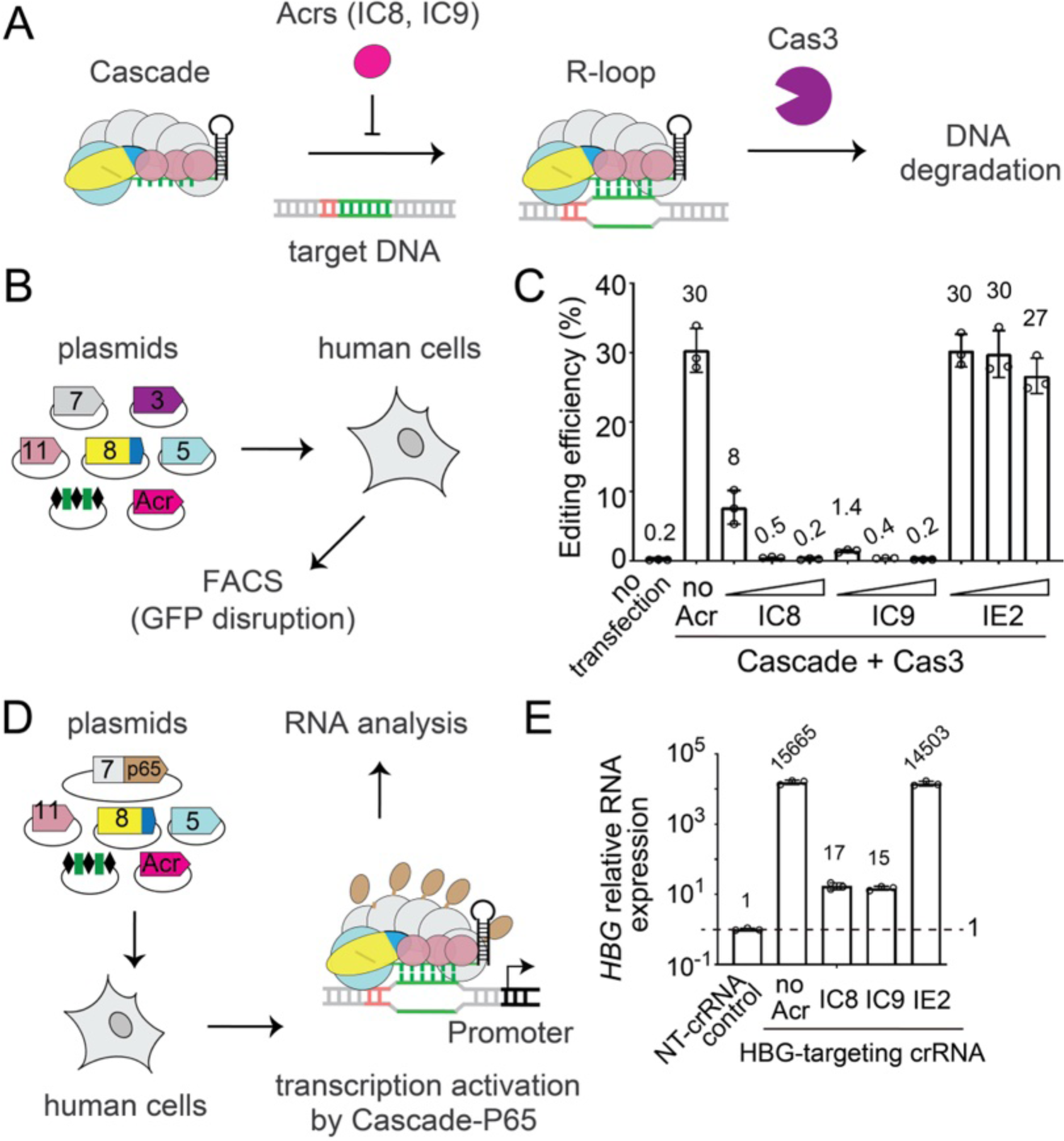
AcrIC8 and IC9 are effective off-switches for Nla I-C CRISPR-Cas3 in human cells. **(A)** Diagram showing AcrIC8 and AcrIC9 could function as off-switches to prevent Nla I-C CRISPR-Cas interference. **(B)** Schematics showing the plasmids used to deliver Acr and Nla I-C components into human cells for RNA-guided GFP knockout. **(C)** FACS sorting based quantification of editing efficiency by Nla I-C Cascade-Cas3 in HAP1-GFP cells, and the inhibitory effect of AcrIC8, AcrIC9, or AcrIC2 at various doses (20, 100, 150 ng of Acr plasmids). 20 ng of AcrIC9 and AcrIC8 plasmid reduced the editing efficiency from 30% to 7.7% and 1.4%, respectively; higher doses reduce editing to the level of no transfection background. AcrIE2 as negative control gave minimal effect. Data shown are mean ± SD, n = 3. **(D)** Schematics showing plasmid-delivery based CRISPR-a in human cells, mediated by the P65 transcription activation domain fused C-terminal to Cas7. Effect of Acr plasmid co-delivery is further evaluated. tuned by Acr plasmid. Cas3 is omitted, Cascade-p65 is directed by crRNA to target *HBG* promoter. **(E)** qPCR quantification of *HBG* RNA expression. Transcription activation is evaluated by normalizing qPCR results against that from the non-targeting control. AcrIC8 and AcrIC9 repressed NlaCascade-p65-mediated *HBG* gene activation by 924- and 1045-fold, respectively, whereas AcrIE2 had no discernible effect. Data shown are mean ± SD, n = 3.

Given that both Acrs block target DNA binding by Cascade (**Figure. 7A**), we also sought to leverage them to control Type I CRISPR-mediated transcriptional modulation. By tethering a P65 transcriptional activator domain to NlaCas7 and omitting Cas3, we established the first Type I-C based endogenous gene activation platform in human cells (**Figure. 7D**). When Cascade-p65 is programmed to target the endogenous *HBG* promoter in HEK293T cells, we observed >15000-fold activation of *HBG* mRNA expression, relative to a non-targeting control Cascade (**Figure. 7E**, first two bars). Importantly, when we co-transfected an additional plasmid encoding human-codon optimized AcrIC8 or AcrIC9 gene, *HBG* transcription was only modestly elevated by 15- and 17-fold, respectively. This result reflected a near one-thousand-fold repression of gene activation by each I-C Acr, whereas the negative control AcrIE2 did not cause discernible inhibition (**Figure. 7E**). In summary, AcrIC8 and AcrIC9 can serve as effective off-switches for eukaryotic genome engineering applications.

## Discussion

While Cascade structures from five subtypes have been reported, their further interactions with the DNA degradation enzyme Cas3 were only defined in I-E and I-A systems (Hu et al., 2022; Xiao et al., 2018). These two examples showcased two distinct mechanistic modes. Cas3 in I-E system interacts with Cascade conditionally, controlled by an R-loop triggered conformational change in Cascade (Xiao et al., 2018). Cas3 in I-A system binds Cascade constitutively; however, its DNase activity is switched on conditionally, only upon R-loop formation (Hu et al., 2022). Existing structural and functional evidence (McBride et al., 2020; Nam et al., 2012; O’Brien et al., 2023; O’Brien et al., 2020; Tan et al., 2022) show that I-C Cascade resembles I-A Cascade in structure but followed the suite of I-E Cascade in Cas3 recruitment and activation mechanisms. Here our I-C Cascade-Cas3 structure rationalizes the mechanistic distinctions in high resolution. The specific Cascade-Cas3 contacts bear no resemblance between I-C and I-E systems, however, the mechanistic principles are shared. Both involve establishing conformationally sensitive molecular contacts to indirectly read out the R-loop state in Cascade. This means the conditional Cas3 recruitment mechanism is under convergent evolutionary selection among multiple subtypes. While most CRISPR systems trigger localized DNA cleavage, Type I CRISPR deletes kilobase pairs of DNA upon activation. The dire consequence likely has led to stronger evolutionary pressure to safeguard off-targeting. The I-C system may represent an evolutionary transition, when the RNA-guided DNA cleavage mechanism in Type I switched from allosteric activation to trans-recruitment and activation, to further reduce the probability of off-targeting.

Anti-CRISPR proteins carry great promises as off-switches to mitigate undesired editing by Cas effectors. It is unlikely to find Acrs that are highly effective against a broad spectrum of CRISPR-Cas systems, as structural variations among Cas homologs may weaken the high-affinity interactions from Acr. Our work underlines the importance of doing systematic screening to identify the most effective Acr against a given genome editor. In Type I systems, high-resolution understanding of Acrs was previously limited to the I-F systems (Chowdhury et al., 2017; Guo et al., 2017; Rollins et al., 2019) **(Figure S8A)**. Together with a recent study (O’Brien et al., 2023), we now have four Acr-bound Cascade structures to expand our mechanistic understanding to I-C systems. Several themes become apparent. All four I-C Acrs bind to the PAM-proximal region of the Cascade, which makes sense as it prevents Cascade from initiating target searching **(Figure S8B-D)**. Except for AcrIC8, the rest three I-C Acrs are highly acidic and mimics dsDNA to various extent. Such Acrs could conceivably serve as broad-spectrum inhibitors. Indeed, AcrIF2/IC2 has been shown to be a versatile inhibitor against multiple Type I subtypes (Csorgo et al., 2020). Interestingly, while effective against Dvu I-C Cascade (O’Brien et al., 2023), AcrIC4 and AcrIC2 posse no inhibitory activity against Nla I-C in our assays. In contrast, AcrIC9 exhibits near-complete inhibition of Nla I-C interference **(Figures 4A-B; S5A-B)**. Our structure rationalizes the high efficacy of AcrIC9, as it not only mimics a dsDNA, but also a dsPAM preferred by Nla Cascade, and locks the PAM-recognition elements in Cascade through highly specific molecular interactions. Unlike others, AcrIC8 is slightly basic in surface charges. We reason that AcrIC8 is highly specific against Nla Cascade because of its ability to establish multifaceted interactions with Nla Cascade. The high-resolution information will inspire new engineering strategies to further strengthen the Type I Acr-Cascade interactions to improve their efficacy as off-switches in the genome editing setting.

### Acr proteins in biotechnological applications

Acrs proteins against the Cas9 or Cas12 nucleases have provided genetically encodable brakes to control gene editing in numerous application contexts (Marino et al., 2020). For instance, delayed introduction of AcrIIA4 after Cas9 RNP delivery can reduce off-target editing in human cells (Shin et al., 2017). Tissue-specific control of Acr expression reinforced safe in vivo Cas9 editing only in the desired organ of mice (Lee et al., 2019). Acrs have also been exploited as safety controls for gene drives, enablers for biosensors and synthetic gene circuits, facilitators for phage therapies, and capture ligands to quantify Cas9 presence in vitro (Kraus and Sontheimer, 2023; Marino et al., 2020). Despite the growing list of Type I CRISPR-enabled eukaryotic applications, no Acr off-switches have been developed to control these technologies. Here we established the first anti-CRISPR inhibitors that can robustly turn off Type I CRISPR activity in human cells. This sets the stage to further improve Type I editing outcomes through temporal, spatial, tissue-specific, and light or drug controlled Acr regulations. The dose-dependent response of AcrIC8 and AcrIC9 **(Figure. 7C)** may allow tunable control beyond a simple on-off switch.

Divergent type I CRISPR editors with guide orthogonality (Tan et al., 2022) can support simultaneous by distinct applications. Narrow spectrum Acrs may allow control of one variant without interfering with the others, whereas broad spectrum Acrs can shield the entire organism from any leaky activity of all editing agents. It is therefore important to understand the cross-reactivity of each Acr against different Type I systems. Lastly, Cas3-specific Acr (e.g., AcrIF3 (Bondy-Denomy et al., 2015; Pawluk et al., 2017) that are known to prohibit Cas3 recruitment to target-bound Cascade have yet to be developed for gene-editing use. They would control Cas3-mediated deletion outcome without impeding Cascade only applications.

## Materials and methods

### Experimental model and subject details

#### *Escherichia coli* BL21(DE3)

*E. coli* BL21 (DE3) were used for protein production. Cells were grown in Lysogeny Broth (LB) supplemented with appropriate antibiotics.

#### *Escherichia coli* BL21-AI

This strain was used for CRISPR interference assay. Cells were grown at 37°C in LB supplemented with appropriate antibiotics and induced with 0.2% L-arabinose and 1 mM IPTG.

#### *Escherichia coli* BW251113 ΔCRISPR-Cas ΔlacIlacZ

A gift from Dr. Chase Beisel. This strain was used for CRISPR repression assay. Cells were grown at 37°C in LB supplemented with appropriate antibiotics and induced with 0.1% L-arabinose.

#### *Escherichia coli* DH5 alpha and JM109

These two strains were used for cloning. Cells were grown at 37 °C in LB supplemented with appropriate antibiotics.

#### HAP1 cell culture

Human HAP1 derivative reporter cell line (HAP1-AAVS-EGFP, Tan *et al*., 2022) were cultured in IMDM (Gibco) supplemented with 10% FBS (Corning) at 37 °C and 5% CO_2_ in a humidified incubator, with daily media change. Cells were split every 2 to 3 days using TrypLE Express (Gibco).

## Method Details

### Purification of Nla Cascade and NlaCas3

Nla Cascade RNP and NlaCas3 protein were each recombinantly expressed in *E. coli* BL21(DE3) cells and purified via Nickel affinity purification followed by SEC, as previously described (Tan *et al*., 2022).

### CRISPR interference assay in *E. coli*

Plasmid interference assays and creation of the BL21-AI derivative strains used were conducted as described (Tan *et al*., 2022) with minor modifications. The pACYC-Duet-based plasmid encodes an extra T7-Acr expression cassette, in addition to the CRISPR array. For strain creation, two plasmids expressing *cas3* and *cascade* were first co-transformed into BL21-AI (Thermofisher). The resulting intermediate strain was made competent using Mix&Go *E. coli* Transformation Kit (Zymo) and transformed with two plasmids, a pCDF1 plasmid containing target sequence and a pACYC-Duet plasmid encoding CRISPR array and Acr, leading to final BL21-AI derivative strains harboring four compatible plasmids. Interference assays were done as described previously, with depletion ratios for both induced and uninduced cultures calculated as the CFUs of triple antibiotic (w/o spectinomycin) control plates divided by CFUs from quadruple-antibiotic test plates.

### CRISPR repression assay in *E. coli*

A plasmid encoding *cascade* and CRISPR array (e.g., pYZ1432) was transformed in *E. coli* strain BW25113 ΔCRISPR-Cas ΔlacIlacZ, a gift (CB414) from Dr. Chase Beisel. The resulting strain was made competent using Mix&Go *E. coli* Transformation Kit (Zymo) and transformed with a PAM-SCANR-derived target plasmid, which contains a Cascade binding site within LacI promoter proceeding the LacI gene. Immediately downstream of lacI is the LacI-dependent lacZ promoter controlling expression of GFP (Leenay et al., 2016). The resulting intermediate strain was made competent and transformed with a 3^rd^ pBAD plasmid encoding *acr* from an arabinose-inducible promoter, leading to a final strain harboring three compatible plasmids.

For CRISPR repression assay, a single colony of this final strain was used to inoculate an overnight culture of 3 mL LB with four antibiotics (20 μg/mL chloramphenicol, 100 μg/mL spectinomycin, 50 μg/mL kanamycin, and 50 μg/mL carbenicillin), with or without 0.1% arabinose as inducer. The cultures were pelleted and resuspended in 1x PBS to OD_600_ of 1 in 1.5 mL Eppendorf tubes, then imaged using a ChemiDoc MP imager (BioRad) on the Alexa488 channel for 0.036 s. These OD_600_-normalized samples were subjected to 2-fold serial dilutions, followed by fluorescence (483/525nm excitation/emission) and absorbance (600 nm) measurements with 96-well black microplates (Corning) and a Safire 2 Multi-detection plate reader (Tecan). The fluorescence value of each sample was normalized by subtracting background fluorescence of the 1x PBS blank control, and then dividing by its OD_600_ number which was also blank subtracted. GFP repression fold (i.e., depletion ratio) was calculated as the normalized fluorescence of arabinose induced samples divided by the normalized fluorescence of uninduced samples.

### Purification of anti-CRISPR proteins

AcrIIA4 protein was purified as described earlier (Rousseau *et al*., 2018). AcrIC8 and AcrIC9 proteins were purified similarly, with minor modifications. Briefly, Acr protein expression was induced with 0.5 mM IPTG at 18 °C for overnight. Cells were pelleted and resuspended in lysis buffer (30 mM HEPES pH 7.5 for AcrIC9 or 30 mM Tris pH 8.5 for AcrIC8, 500mM NaCl, 20 mM imidazole, 0.5 mM TCEP, 1x FastBreak lysis reagent) and lysed by sonication. Clarified lysate was purified with Ni-NTA resin and His-tagged Acr proteins were eluted in elution buffer (30 mM HEPES pH 7.5 for AcrIC9 or 30 mM Tris pH 8.5 for AcrIC8, 500 mM NaCl, and 300 mM imidazole). The eluate was mixed with His-tagged TEV protease for AcrI9 or His-tagged 3C protease for AcrIC8, and dialyzed (30 mM HEPES pH 7.5 for AcrIC9 or 30 mM Tris pH 8.5 for AcrIC8, 300 mM NaCl, and 0.5 mM DTT) overnight at 4°C. Cleaved, untagged Acr proteins were purified using new Ni-NTA resin by collecting the unbound fraction, concentrated, and further purified by loading onto a sephacryl S200 column using the same dialysis buffer as SEC elution buffer.

### Acr-Cascade nickel pull-down

Untagged 3.75 μM anti-CRISPR proteins were incubated with or without 750 nM His-tagged Cascade or His-tagged Cas3 on ice for 30 min in 1x binding buffer (30 mM HEPES pH 7.5, 300 mM NaCl, 0.5 mM DTT). As negative controls, His-tagged Cascade or His-tagged Cas3 was incubated on their own without any Acr. After incubation, input fractions were set aside for SDS-PAGE analysis and MagneHis Ni Particles (Promega) were added to each reaction. After 30 min incubation at 4°C with agitation, the tubes were placed on the magnetic stand to immobilize the beads at tube wall and the supernatant discarded. The beads were washed 3 times with 500 μL binding buffer, collected on the magnetic rack, and resuspended in 20 μL binding buffer. Beads and input fractions were diluted 1:2 with 2x Laemmli buffer (Bio-Rad) supplemented with 100 mM DTT and analyzed by 15% SDS-PAGE followed by Coomassie blue staining.

### Electrophoretic mobility shift assay (EMSA)

*In vitro* binding was conducted in 1x binding buffer (30 mM HEPES pH 7.5, 150 mM NaCl). Cascade RNP was diluted to the indicated concentrations (3, 6, 12, 25, 50, 100 nM). Fluorescently labeled dsDNA substrate was then added (16 nM). The binding reactions were incubated at 37°C for 30 min, 10% (v/v) glycerol was added, and samples resolved by 1% agarose/1X TAE gel electrophoresis. Gels were by imaged with a ChemiDoc MP imager (Biorad). Assays were conducted in triplicate with a representative gel shown.

### Plasmid transfection into human cells and flow cytometry

CRISPR-Cas and Acr plasmid mixture was transfected into HAP1-EGFP reporter cells and editing efficiency measured by flow cytometry as previously described (Tan et al., 2022), with minor modifications. Cells were seeded one day before transfection at 0.6×10^5^ cells per well of a 24-well plate. For each transfection, we used a total of 350 ng *crispr-cas* plasmids (27, 13.5, 40.5, 162, 27, and 30 ng of *cas3*, *cas5*, *cas7*, *cas8*, *cas11* and *crispr* plasmids, respectively). Acr-encoding plasmids were tested at 20, 100, and 150 ng. Flow cytometry data were analyzed with FlowJo v10.8.1.

### Cascade-mediated Cas3 DNA cleavage assay

The 192 bp dsDNA substrate was produced from PCR using 5′-fluorescently labeled primers (FAM at NTS and Cy5 at TS). The reaction mixture was prepared from 100 nM final concentration of Cascade, 100 nM Cas3 and 10 nM substrate in a cleavage buffer containing 10 mM HEPES pH 7.5, 150 mM NaCl, 10 mM MgCl2 with or without 2 mM ATP. The reaction was incubated at 30°C for 30 min. After incubation, the nucleic acids were phenol-chloroform extracted and separated on a 12% denaturing urea polyacrylamide gel. Fluorescent signals were recorded using a Typhoon™ scanner (Amersham).

### Cryo-EM sample preparation

250 μL Cascade at 5 μM concentration was incubated with 50 μL the specified target RNA at 50 μM in binding buffer (150 mM NaCl, 25 mM HEPES pH 7.5) for 15 minutes at room temperature. Then added 50 μL Cas3 at 50 μM and incubated for 15 minutes. Then loaded the complex on superdex 200 column. Pooled the complex peak and concentrated to 0.5 mg/ml concentration. 3.5 μL of the final complex was applied to a Quantifoil holey carbon grid (1.2/1.3, 200 mesh) which had been glow discharged with 20 mA at 0.39 mBar for 30 seconds (PELCO easiGlow). Grids were blotted with Vitrobot blotting paper (Ted Pella Inc STANDARD VITROBOT FILTER PAPER; Cat# 47000100) for 3 seconds at 4 °C, 100% humidity, and plunge-frozen in liquid ethane using a Mark IV FEI/Thermo Fisher Vitrobot.

For the cryo-EM data collection of Cascade-Acrs, 25 μL Cascade at 2 μM was mixed with 5 ul Acrs at 50 μM in the binding buffer and incubated for 30 min on ice. 3.5 μL of the final complex was applied to a Quantifoil holey carbon grid (1.2/1.3, 200 mesh) which had been glow discharged with 20 mA at 0.39 mBar for 30 seconds (PELCO easiGlow). Grids were blotted with Vitrobot blotting paper (Ted Pella Inc STANDARD VITROBOT FILTER PAPER; Cat# 47000100) for 5 seconds at 4 °C, 100% humidity, and plunge-frozen in liquid ethane using a Mark IV FEI/Thermo Fisher Vitrobot.

### EM data acquisition

All data was collected on a 200 kV Talos Arctica (Thermo Scientific) with Gatan K3 direct electron detector. The total exposure time of each movie stack led to a total accumulated dose of 50 electrons per Å2 which fractionated into 50 frames without beam tilt. Dose-fractionated super-resolution movie stacks collected from the Gatan K3 direct electron detector were binned to a pixel size of 1.1 Å. The defocus value was set between −1.0 μm to −2.5 μm.

### Image processing and 3D reconstruction

Motion correction, CTF-estimation, blob particle picking, 2D classification, 3D classification and non-uniform 3D refinement were performed in CryoSPARC v.2. Refinements followed a standard procedure, 100 classes of of 2D classification was performed in CryoSPARC. High quality 2D classes were hand-picked to do 3D classifications in C1 symmetry. CryoSPARC was instructed to generate eight initial models in 3D classification for each reconstruction. After heterologous refinement, the promising 3D models was hand-picked for post 3D refinement, as outlined in the supplemental figures. A solvent mask was generated in RELION with 0.1 contour level and was used for all subsequent local refinement steps. CTF post refinement was conducted to refine the beam-induced motion of the particle set, resulting in the final maps. The detailed data processing and refinement statistics for cryo-EM structures are summarized in supplemental figures and **Table S2**.

### Atomic model building

AlphaFold2 prediction was individually docked into the cryo-EM map using UCSF Chimera. Model building was completed through iterative cycles of manual building using Coot and real space and positional refinements using Phenix. The detailed statistics are documented in **Table S2**.

### Model Validation

Map-model Fourier Shell Correlation (FSC) was computed using Mtriage in Phenix. Map-model FSC resolution of each dataset was estimated from Mtriage FSC curve, using 0.143 cutoff. The masked cross-correlation (CCmask) from Mtriage was reported as representative model-map cross-correlation. Model geometry was validated using the MolProbity. All the deposited models were submitted to MolProbity server to check the clashes between atoms, Ramachandran-plot, bond-angles, bond-lengths, sidechain rotamers, CaBLAM and C-beta outliers. All the model validation stats are summarized in **Table S2**.

## Acknowledgments

This work is supported by the National Institutes of Health grants GM118174 to A.K. and GM137883 to Y.Z., and the University of Michigan Biological Scholar Award to Y.Z. M.T.M. was supported by the NIH T32 Predoctoral Training grant in Genetics T32GM007544. M.T.M and M.L.L. are each supported by a University of Michigan Rackham Graduate Student Research Grant. This work made use of facilities supported by the National Science Foundation Materials Research Science and Engineering Centers program grant DMR-1719875 (Cornell Center for Materials Research Shared DOE Office of Biological and Environmental Research grant KP1607011 (Laboratory for BioMolecular Structure [LBMS]). We thank Jeremy W. Dortch and Jue Chen for technical assistance and helpful discussions. We thank Chase L. Beisel for sharing plasmids and *E. coli* strains to develop the GFP repression assay.

## Author contributions

C.H., M.T.M., Y.Z. and A.K. designed the research. C.H. is responsible for the cryo-EM reconstructions and cleavage assays. X.Z. and Z.H. purified all proteins. M.T.M. performed EMSA, His pull-down, and *E. coli* interference and CRISPRi assays. M.L.L. and X.Z. carried out initial cloning and interference test of all Acrs. M.T.M conducted all human cell work. A.K. Y.Z. C.H. and M.T.M wrote the manuscript with input from Z.H. and X.Z.

---

**Figure S1.**
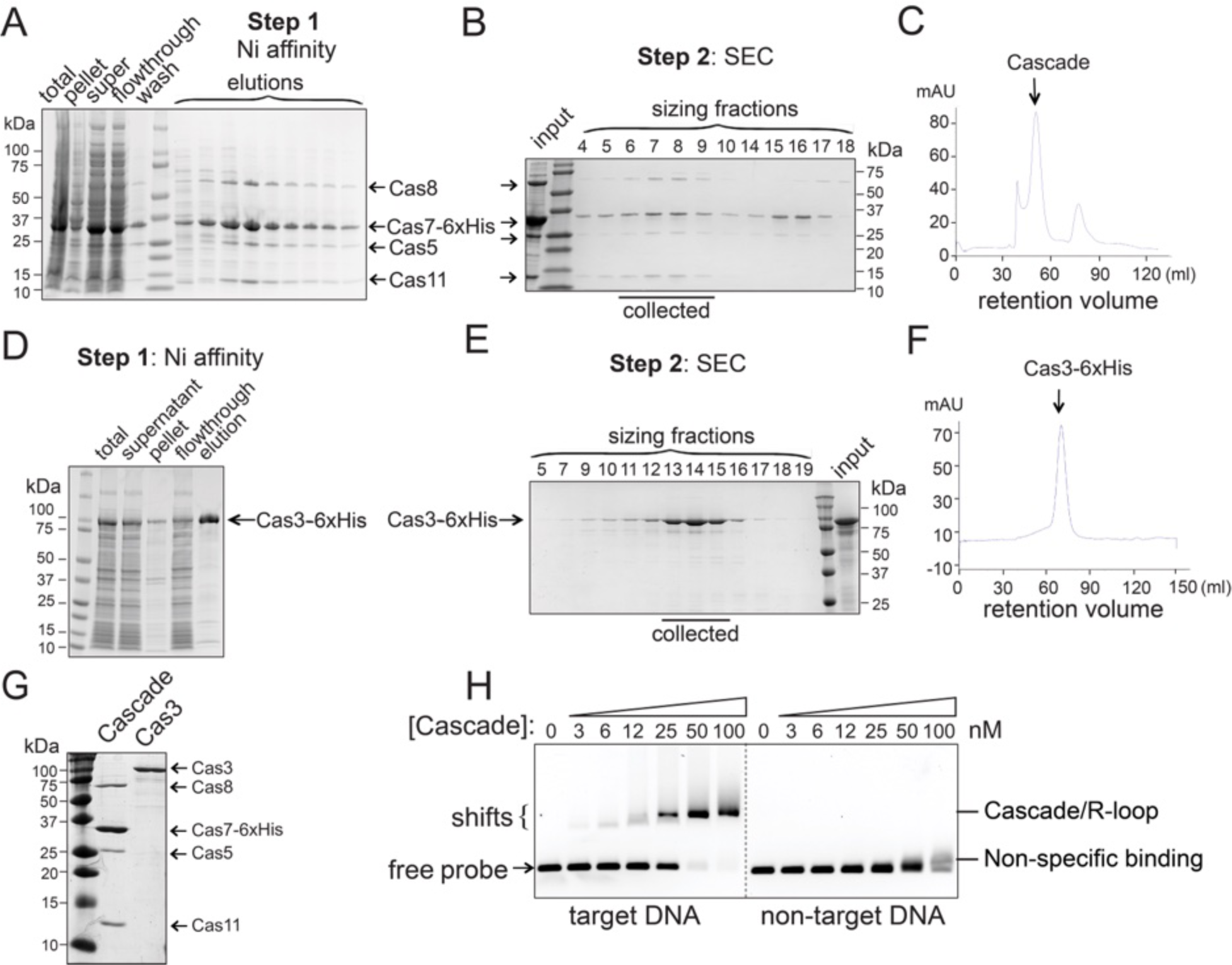
Reconstitution and characterization of Nla CRISPR-Cas3, related to Figure 1. **(A)** Nla Cascade complex was reconstituted in *E. coli* and purified on Ni-NTA affinity column via a C-terminal His_6_ tag on Cas7, followed by size-exclusion chromatography (SEC). **(B, C)** SEC fractions evaluated on SDS-PAGE and its elution profile, respectively. **(D)** SDS-PAGE of Nla Cas3 purification via a C-terminal His_6_-tag. **(E, F)** SDS-PAGE and elution profiles from the second step SEC purification, respectively. **(G)** Quantity of the Nla Cascade and Cas3 used in cryo-EM and biochemical assays, analyzed using Coomassie-blue stained SDS-PAGE. **(H)** EMSAs showing robust RNA-guided DNA binding by Nla Cascade. Purified Cascade complex (0-100 nM) was incubated with fluorescently labeled dsDNA substrates (16 nM) for 30 min at 37°C, reaction mixtures were then resolved by 1% native agarose gel.

**Figure S2.**
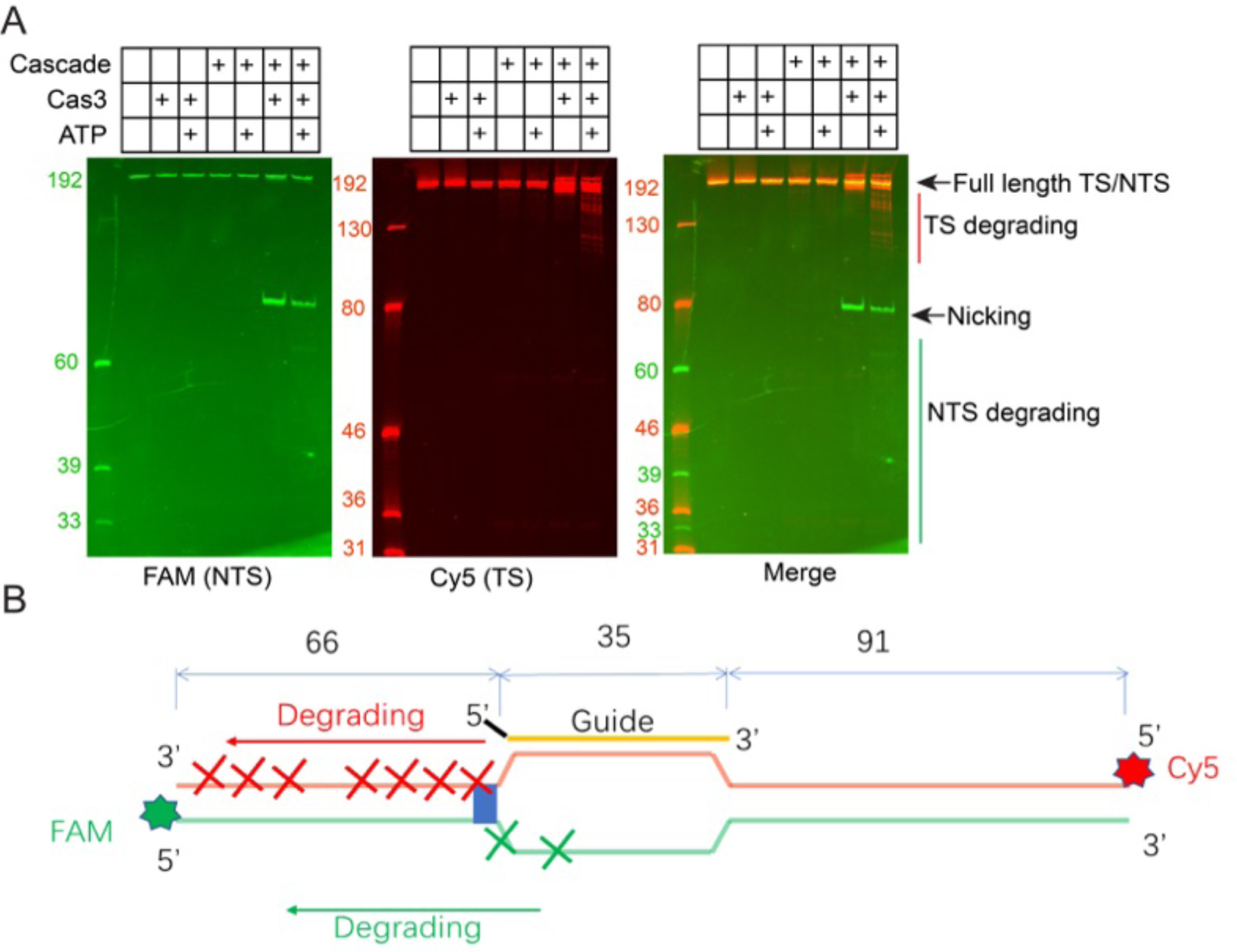
Nla Cascade-Cas3 mediated target DNA cleavage *in vitro,* Related to Figure 1. **(A)** Representative urea-PAGE gel image resolving the DNA cleavage products by Nla I-C Cascade-Cas3. The same gel was scanned at FAM (left) and Cy5 (middle) emission wavelength to resolve the cleavage pattern on NTS and TS strands, respectively. The merged image is displayed to the right. **(B)** A sketch depicting the cleavage pattern by Cascade-Cas3 in the presence of ATP.

**Figure S3.**
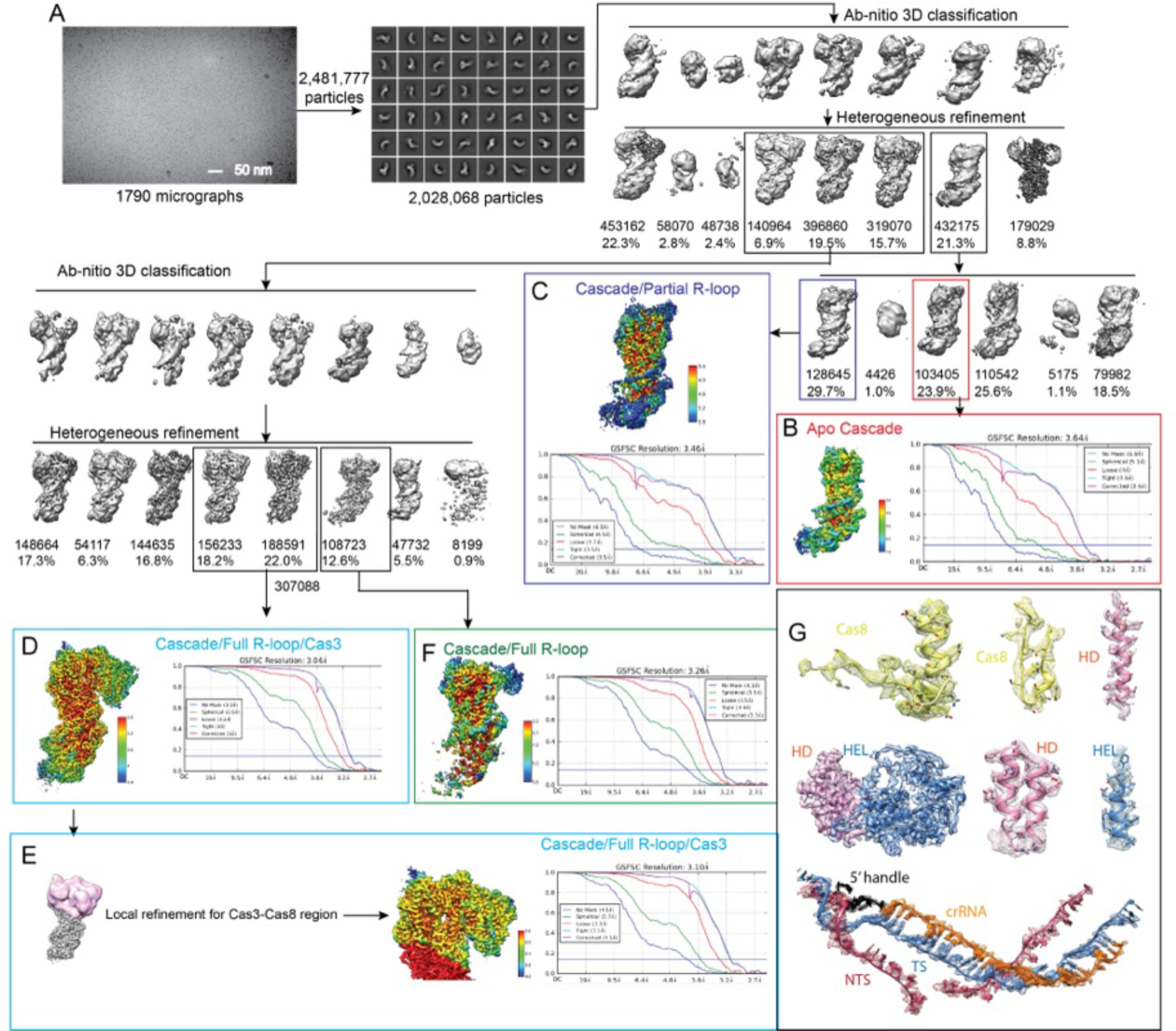
Cryo-EM single particle reconstructions procedure to resolve apo, partial R-loop, full R-loop, and Cas3-bound R-loop states from the DNA-bound Nla Cascade-Cas3 sample, related to Figure 1. **(A)** Cryo-EM image processing and 3D reconstruction procedure for the DNA-bound Nla Cascade-Cas3 sample. We resolved four functional states from this sample: Apo Cascade **(B)**, Cascade/Partial R-loop **(C)**, Cascade/Full R-loop/Cas3 **(D)**, its masked, focused local refinement **(E)**, and Cascade/Full R-loop without Cas3 **(F)**. **(G)** Representative cryo-EM densities showing the quality of the Cascade-Cas3/R-loop reconstruction.

**Figure S4.**
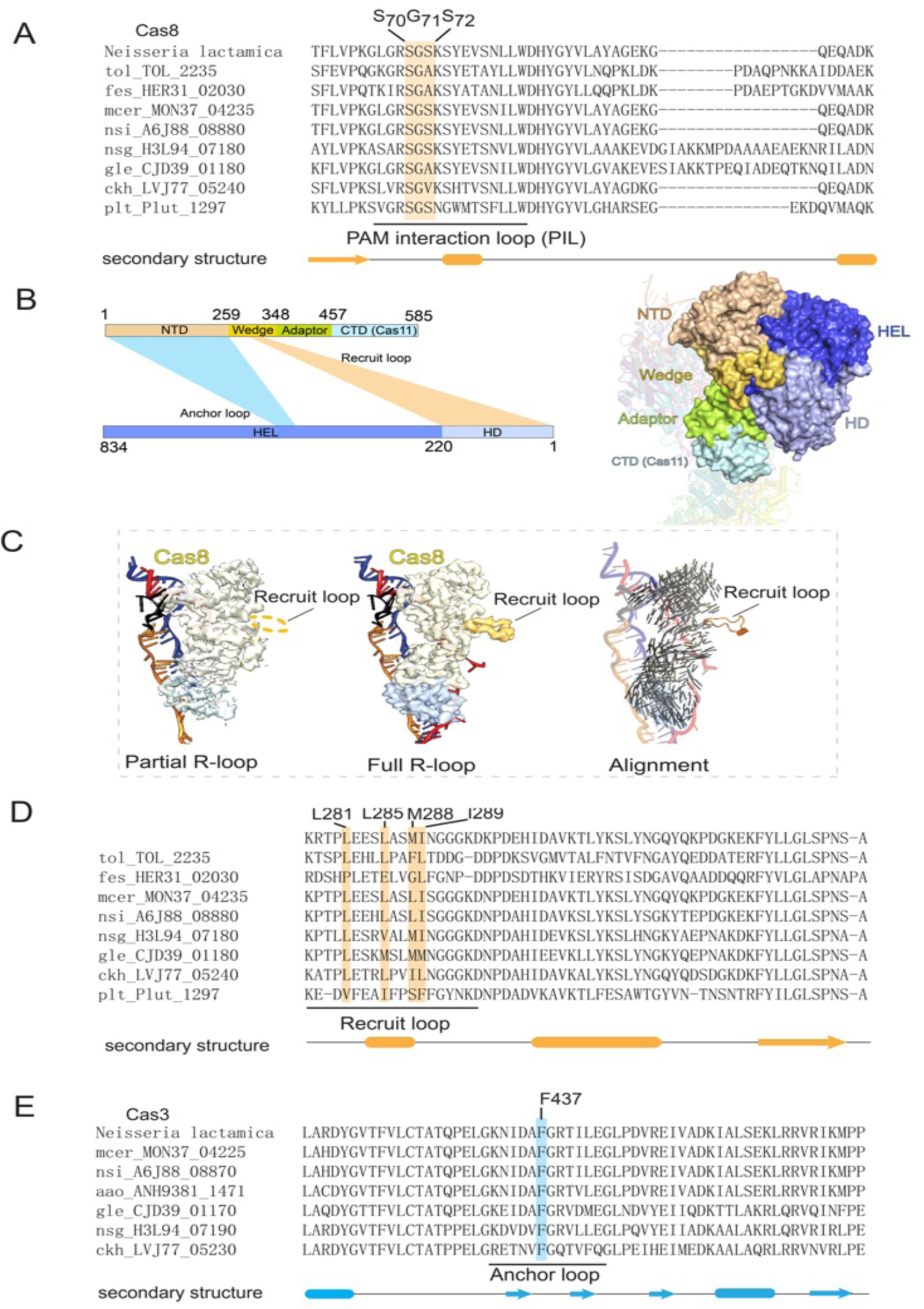
In-depth analyses of PAM recognition and Cascade-Cas3 interactions, related to Figures 2 and 3. **(A)** Multiple sequence alignment (MSA) of Cas8 homologs reveals the conservation of the SGS-motif in the PAM recognition loop. **(B)** Left: the sketch depicting the location of the interactions in Cas8 (top) and Cas3 (bottom). Right: The same interaction in 3D. **(C)** Cas8 conformational changes between partial and full R-loop states. Short bars (right panel) depict the direction and distance of the local movement. **(D)** Cas8 MSA revealing the Recruit loop residues are conserved and hydrophobic. **(E)** Cas3 MSA revealing the tip of the Anchor loop consists of a conserved phenylalanine residue.

**Figure S5.**
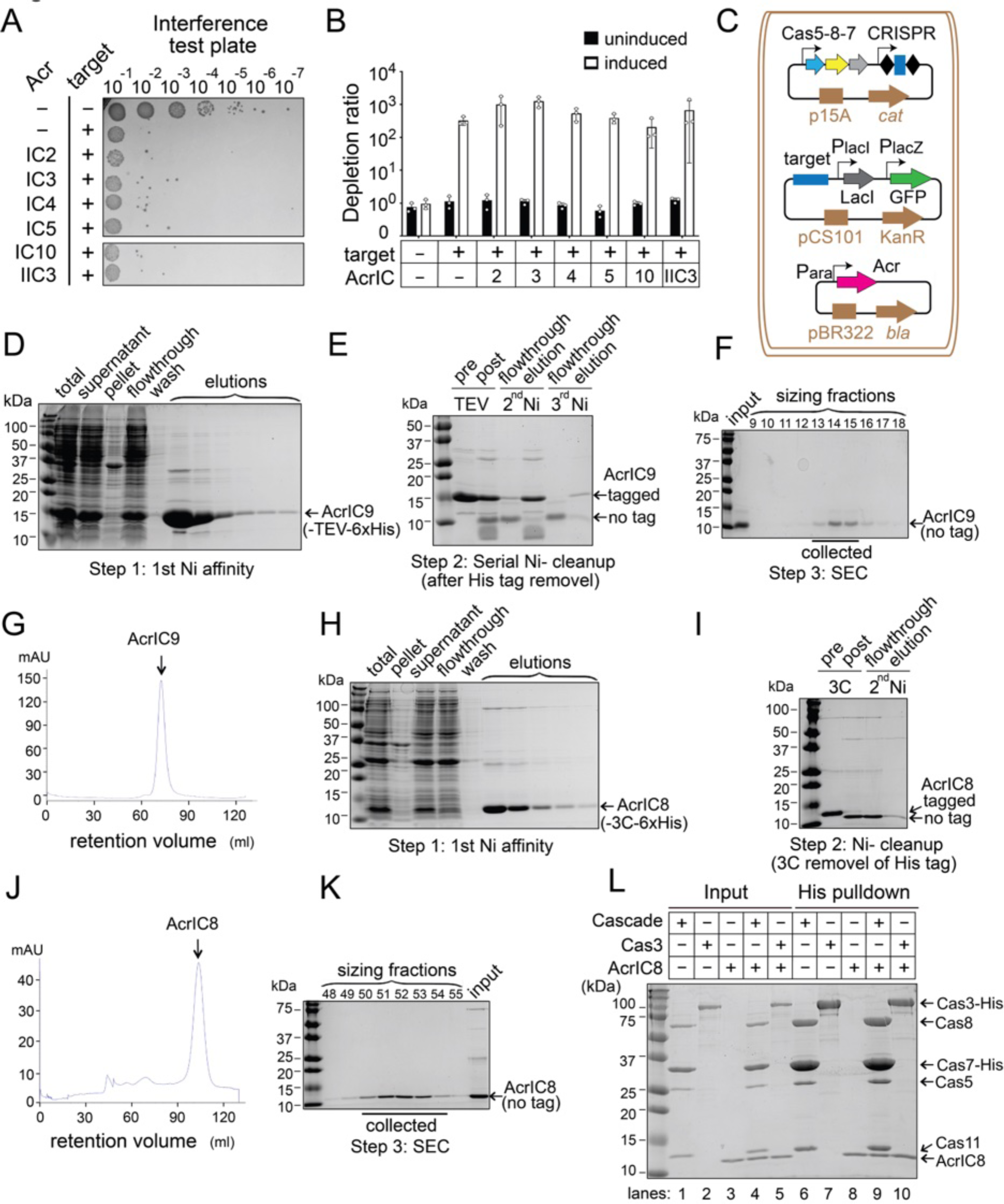
Identification of effective Acr proteins of Nla CRISPR-Cas3 system and biochemical analyses, related to Figure 4. **(A-B)** Acrs IC2, IC3, IC4, and C10 cannot inhibit plasmid interference mediated by Nla I-C CRISPR-Cas. Representative images of *E. coli* interference assay are in (A), where *E. coli* strains each encoding *crispr-cas* genes, a matching functional target, and a candidate Acr gene were spot-titrated onto quadruple-antibiotic test plates. Depletion ratios plotted in (B) are quantified as in Fig. 2E. Data shown are mean ± SD, n = 3. The *acrIC1*-expressing *E. coli* strain failed to grow properly and therefore is not included in this interference test. **(C)** A schematic of the *E. coli* BW25113 derivative strains used in CRISPR repression assay, all harboring three compatible plasmids. The 1^st^ plasmid constitutively expresses *cascade* operon from J23119 promoter and CRISPR array from its native promoter. The 2^nd^ target plasmid drives GFP reporter expression from a LacZ promoter, which can be repressed by LacI repressors produced from an upstream LacI cassette. NlaCascade binding to its target site embedded in the LacI promoter will repress LacI production and thereby re-activates GFP expression. The 3^rd^ plasmid carries an *acr* gene under control of an arabinose-inducible promoter. Induction of an Acr that can prevent NlaCascade from binding to its DNA target will relieve the transcriptional repression of LacI promoter and thus block GFP activation. **(D-G)** Purification of untagged AcrIC9 protein. Recombinant His_6_-TEV-AcrIC9 was purified from *E. coli* first by Ni-NTA affinity pulldown (D), followed by two iterative rounds of TEV protease cleavage & Ni-NTA cleanup for tag removal(E), and a final SEC step (F) with FPLC elution profile showing the AcrIC9 peak in (G). **(H-K)** Purification of untagged AcrIC8 protein. His_6_-3C-AcrIC8 was purified from *E. coli* first by Ni-NTA affinity chromatography (H), followed by 3C protease cleavage & Ni-NTA cleanup to remove epitope tag (I) and a final SEC step (J) with FPLC elution profile showing the AcrIC8 peak (K). **(L)** Untagged AcrIC8 protein alone sticks to nickel affinity resin (lane 8) used in the pulldown experiments, thus precluding any conclusions about its *in vitro* binding capacity to His_6_-Cascade or His_6_-Cas3. The nickel affinity pulldown assay was carried out and analyzed as in Fig. 4G. The input and pulldown fractions are on the left and right of this Coomassie-stained SDS-PAGE gel, respectively.

**Figure S6.**
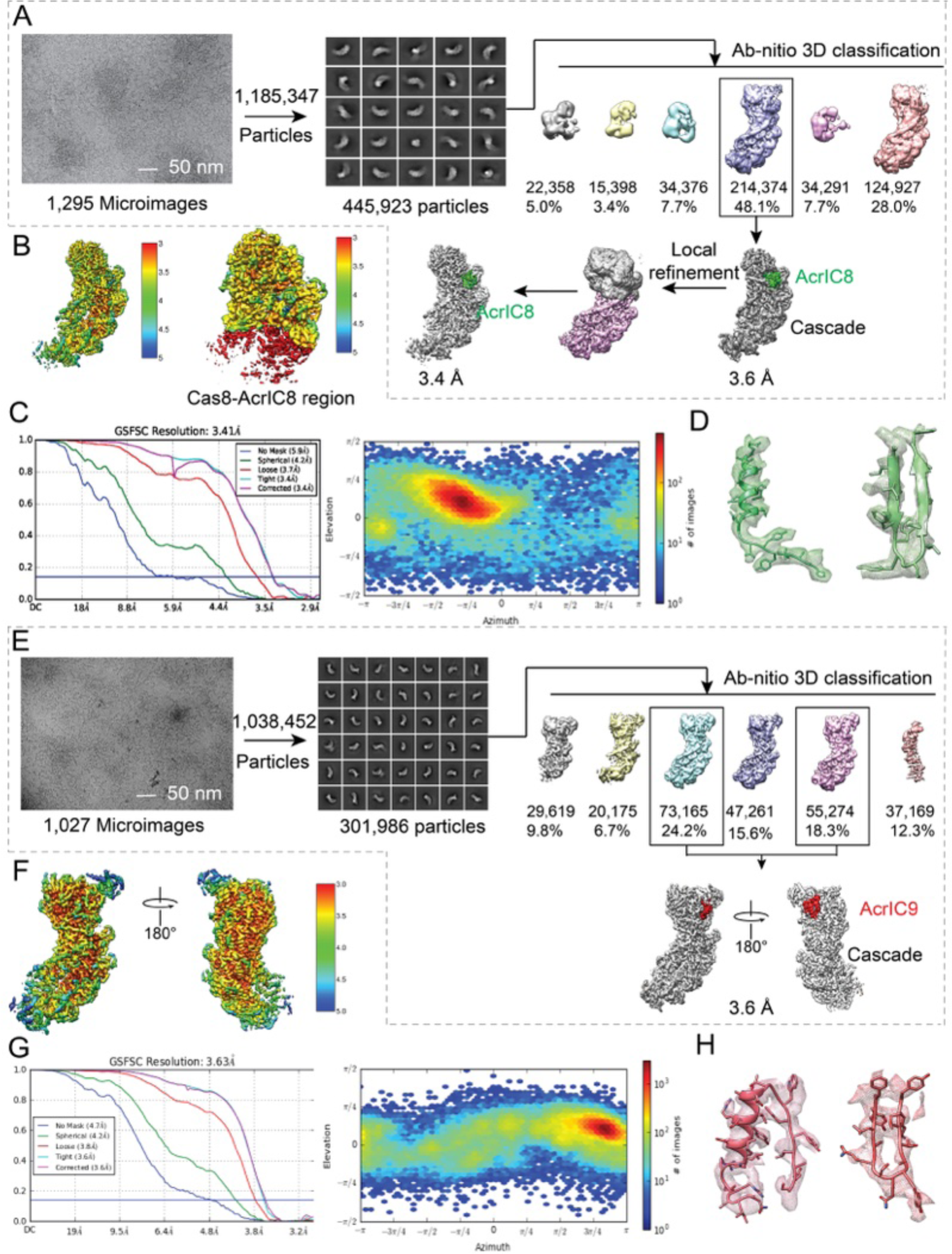
Flow-chart of the cryo-EM single particle reconstructions of Nla Cascade in complex with AcrIC8 and AcrIC9, related to Figures 5 and 6. **(A)** Cryo-EM image processing and 3D reconstruction procedure for the AcrIC8-bound Nla Cascade. **(B)** Local density and resolution. **(C)** Left: the GSFSC chart displays the map resolution. Right: the 2D plot distribution displays the direction distribution. **(D)** Representative AcrIC8 densities. **(E)** Cryo-EM image processing and 3D reconstruction procedure for the AcrIC9-bound Nla Cascade. **(F)** Local density and resolution. **(G)** Left: the GSFSC chart displays the map resolution. Right: the 2D plot distribution displays the direction distribution. **(H)** Representative AcrIC9 densities.

**Figure S7.**
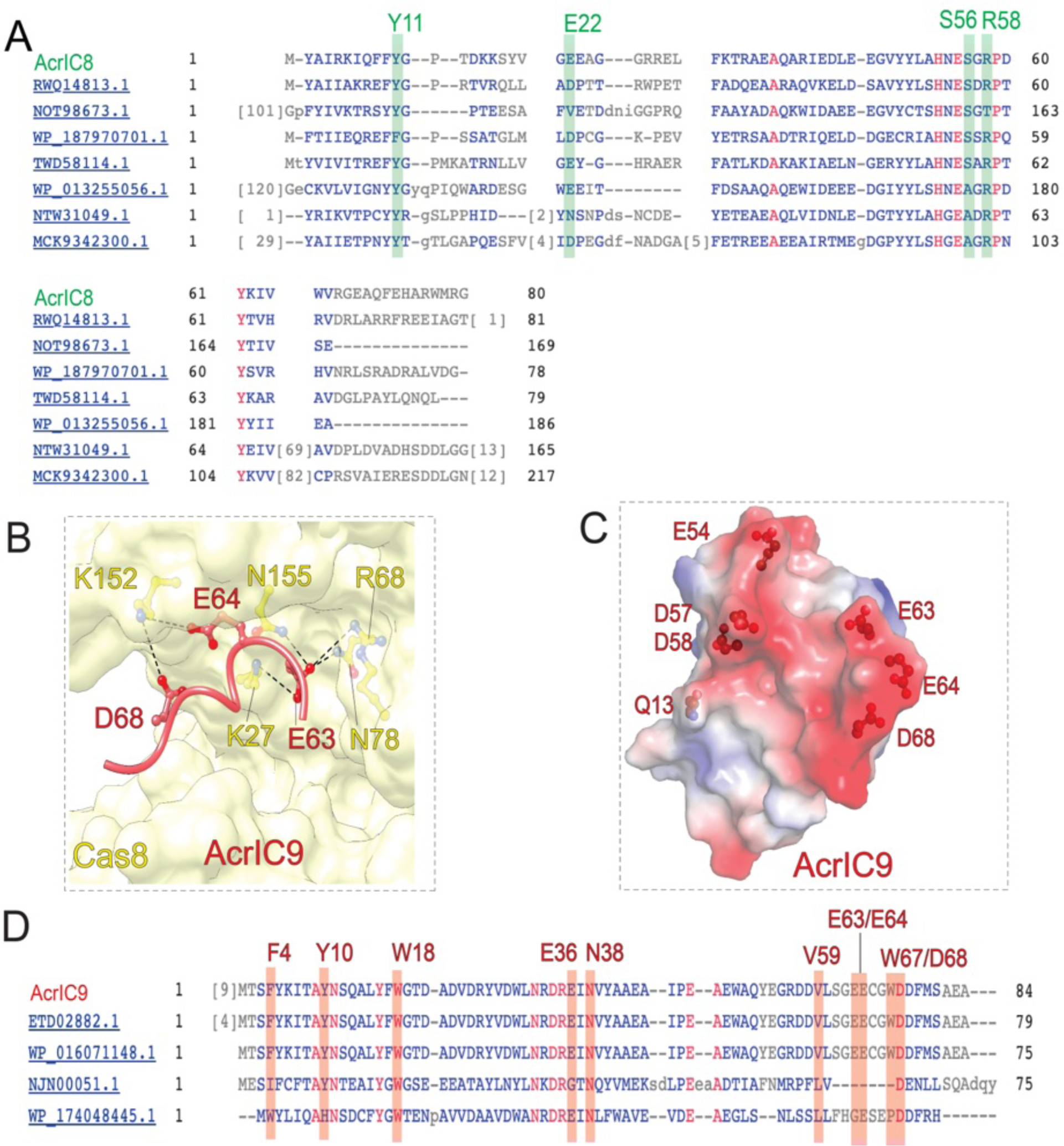
In-depth analyses of the Cascade inactivation mechanisms by AcrIC8 and AcrIC9, related to Figures 5 and 6. **(A)** MSA among AcrIC8 homologs revealing the residue conservation at the Cascade-interacting surface. **(B)** 3D structure showing an acidic patch in AcrIC9 mimics the sugarphosphate backbone of DNA and makes favorable electrostatic contacts with Cas8. **(C)** The electrostatics surface of AcrIC9 showing two parallel stretches of negative charges, mimicking a double-stranded DNA. **(D)** MSA among AcrIC9 homologs revealing sequence conservation at the Cascade-contacting surface.

**Figure S8.**
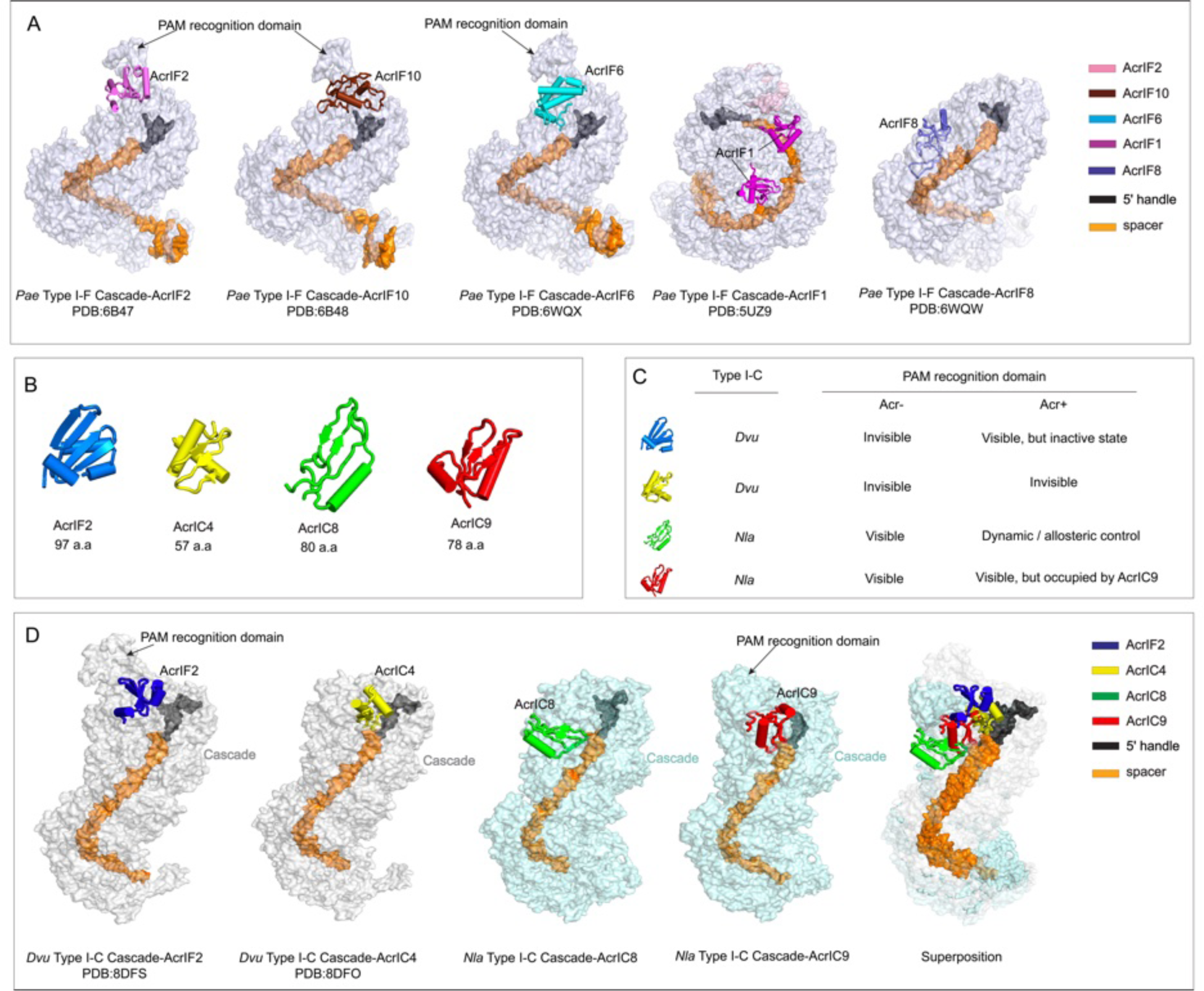
In-depth analyses of the inactivation mechanism of Acr proteins in other type I CRISPR-Cas3 systems, related to Figures 5 and 6. **(A)** Summarization of structures containing Acr-bound Type I-F Cascades. AcrIF2, AcrIF10 and AcrIF6 block DNA targeting by occupying the PAM recognition domain, and AcrIF1 and AcrIF8 block DNA targeting by preventing base-pairing between crRNA and target DNA. **(B)** Structural comparison revealing the wide range of structural diversity among AcrIF2, AcrIC4, AcrIC8 and AcrIC9. **(C)** Summary of the inactivation mechanisms by AcrIF2, AcrIC4, AcrIC8, and AcrIC9. **(D)** Side-by-side comparison of Acrs targeting I-C Cascade. The left two structures targets Dvu Cascade, the middle two structures target Nla Cascade. The right-most structure depicts four Acrs superimposed onto the I-C Cascade, highlighting their different binding locations on Cascade.

**Table S1.**
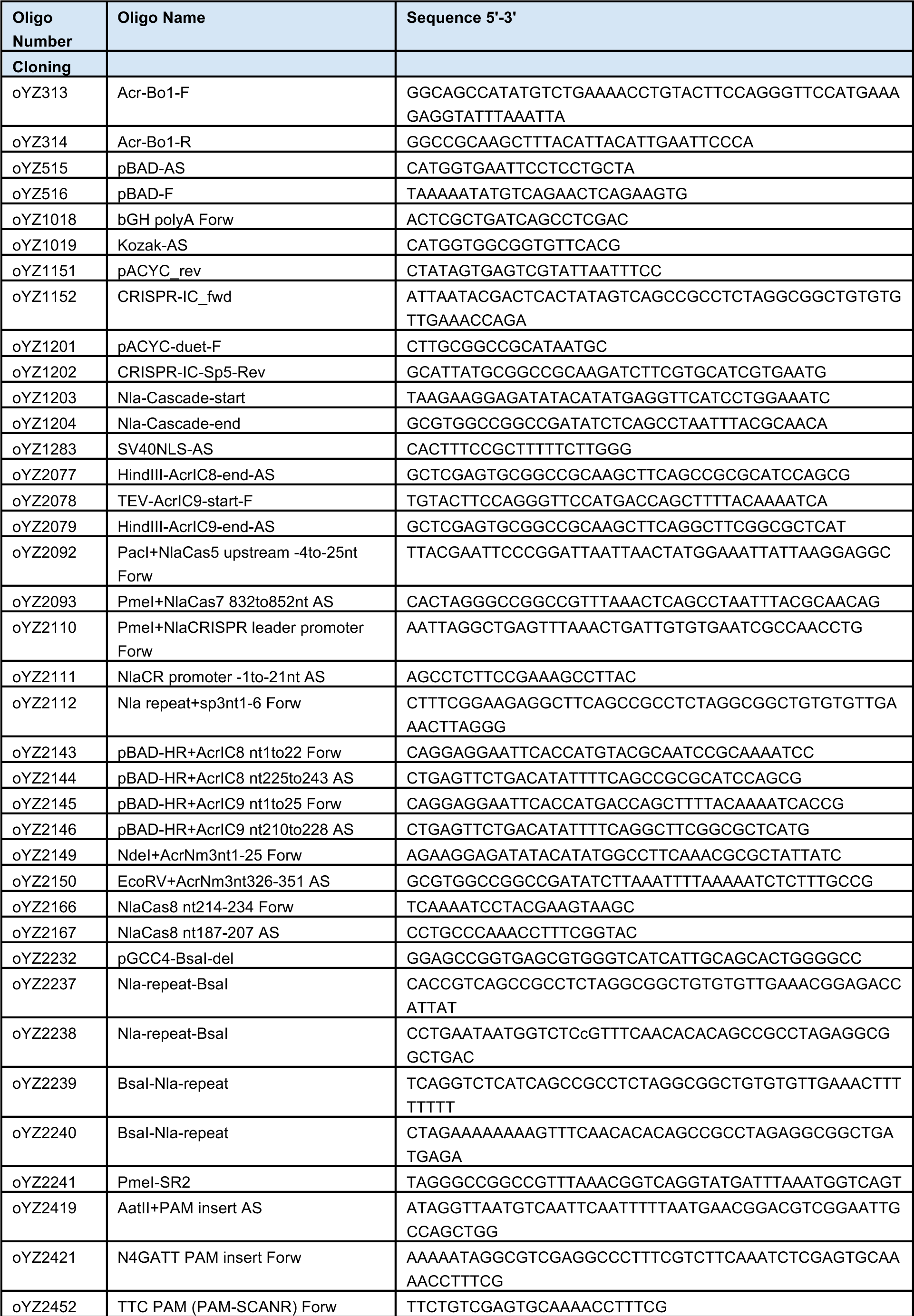
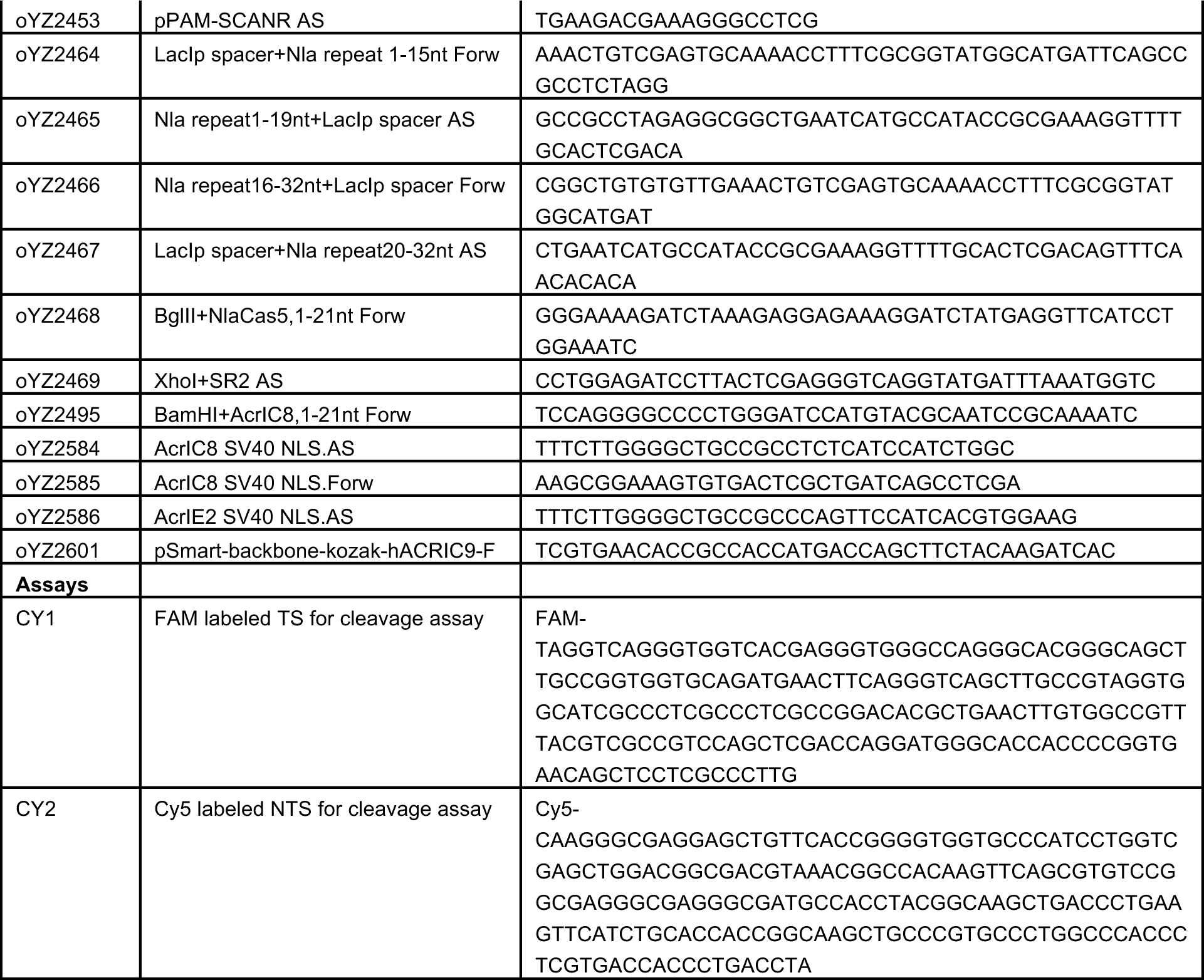
Documentation of oligos, plasmids, and strains used in the study.

**Table S2.** Cryo-EM data collection, refinement and validation statistics.

| Data collection and processing | Cascade alone | Cascade/Partial R-loop | Cascade/Full R-loop | Cascade/Full R-loop/Cas3 | AcrlC8-Cascade | AcrlC9-Cascade |
| --- | --- | --- | --- | --- | --- | --- |
| PDB | 8GAF | 8GAM | 8GAN | 8G9U | 8G9S | 8G9T |
| EMD | 29896 | 29900 | 29901 | 29879 | 29877 | 29878 |
| Magnification(nominal) | 67k |  |  |  | 67k | 67k |
| Voltage (kV) | 200 |  |  |  | 200 | 200 |
| Electron exposure(e-/Å<sup>2</sup>) | 50 |  |  |  | 50 | 50 |
| Defocus range(um) | 1.-2.5 |  |  |  | 0.8-2.5 | 0.8-2.5 |
| Raw Pixel size(Å) | 0.615 |  |  |  | 0.65 | 0.615 |
| symmetry | C1 |  |  |  | C1 | C1 |
| Micrographs | 1,790 |  |  |  | 1,295 | 1,027 |
| Initial particle (No.) | 2,028,068 |  |  |  | 445,923 | 301,986 |
| Final particle (No.) | 103,405 | 128,645 | 108,723 | 307,088 | 214,374 | 128,439 |
| Map resolution(Å) | 3.64 | 3.46 | 3.26 | 3.04 | 3.41 | 3.63 |
| FSC threshold | 0.143 | 0.143 | 0.143 | 0.143 | 0.143 | 0.143 |
| Map resolution range (Å) | 20-3.0 | 20-2.9 | 20-2.8 | 20-2.2 | 20-2.8 | 20-2.8 |
| Refinement |  |  |  |  |  |  |
| Initial model used | AlphaFold2 | AlphaFold2 | AlphaFold2 | AlphaFold2 | AlphaFold2 | AlphaFold2 |
| Map sharpening B factor(Å<sup>2</sup>) | -100 | -100 | -100 | -100 | -103 | -100 |
| **Model composition** |  |  |  |  |  |  |
| Nonhydrogen atoms | 24582 | 25360 | 27669 | 33546 | 25758 | 26804 |
| Protein residues / Nucleotide | 2931/43 | 2961/81 | 3109/136 | 3866/136 | 3102/42 | 3184/43 |
| Ligands | - | - | - | 6 | - | - |
| Water | - | - | - | - | - | - |
| **B factors (Å<sup>2</sup>)** |  |  |  |  |  |  |
| Protein (min/max/mean) | 50.7/500.3/147.2 | 45.6/231.4/91.5 | 59.9/214.4/118.7 | 23.2/121.7/56.2 | 74.4/481.4/173.1 | 47.2/224.5/96.3 |
| Nucleotide (min/max/mean) | 69.5/244.8/114.6 | 56.1/175.2/93.8 | 72.8/224.7/129.2 | 29.8/175.2/59.1 | 99.3/231.6/125.8 | 56.8/150.5/83.0 |
| Ligand | --- | --- | --- |  | --- | --- |
| Water | --- | --- | --- | --- | --- | --- |
| R.m.s.d deviations | 0.00 | 0.00 | 0.00 | 0.03 | 0.00 | 0.00 |
| Bond lengths (Å) | 0.001 | 0.013 | 0.005 | 0.011 | 0.008 | 0.013 |
| bond angles(°) | 0.392 | 1.384 | 0.913 | 1.067 | 0.784 | 1.698 |
| **Validation** |  |  |  |  |  |  |
| MolProbity score | 2.12 | 2.90 | 2.69 | 2.32 | 2.15 | 2.62 |
| Clashscore | 10.53 | 11.68 | 15.21 | 41.6 | 24.07 | 50.52 |
| Poor rotamers(%) | 0.00 | 0.00 | 3.01 | 0.55 | 0.00 | 3.5 |
| Ramachandran plot |  |  |  |  |  |  |
| Favored (%) | 82.67% | 83.32% | 86.01% | 82.96 % | 91.44 % | 89.42 % |
| Allowed (%) | 14.40% | 16.51% | 13.73% | 16.7% | 8.52 % | 10.49 % |
| Disallowed (%) | 2.60% | 0.17 | 0.26% | 0.34% | 0.03 % | 0.09 % |

